# Protective pan-ebolavirus combination therapy by two multifunctional human antibodies

**DOI:** 10.1101/2021.05.02.442324

**Authors:** Pavlo Gilchuk, Charles D. Murin, Robert W. Cross, Philipp A. Ilinykh, Kai Huang, Natalia Kuzmina, Viktoriya Borisevich, Krystle N. Agans, Joan B. Geisbert, Robert H. Carnahan, Rachel S. Nargi, Rachel E Sutton, Naveenchandra Suryadevara, Seth J. Zost, Robin G. Bombardi, Alexander Bukreyev, Thomas W. Geisbert, Andrew B. Ward, James E. Crowe

## Abstract

Ebolaviruses cause a severe and often fatal illness with the potential for global spread. Monoclonal antibody-based treatments that have become available recently have a narrow therapeutic spectrum and are ineffective against ebolaviruses other than Ebola virus (EBOV), including medically important Bundibugyo (BDBV) and Sudan (SUDV) viruses. Here we report the development of a therapeutic cocktail comprising two broadly neutralizing human antibodies rEBOV-515 and rEBOV-442 that recognize non-overlapping sites on the ebolavirus glycoprotein (GP). Antibodies in the cocktail exhibited synergistic neutralizing activity and resisted viral escape, and they were optimized for their Fc-mediated effector function activities. The cocktail protected non-human primates from ebolavirus disease caused by EBOV, BDBV, or SUDV with high therapeutic effectiveness. High-resolution structures of the cocktail antibodies in complex with GP revealed the molecular determinants for neutralization breadth and potency. This study provides advanced preclinical data to support clinical development of this cocktail for pan-ebolavirus therapy.

## Introduction

The Filoviridae family consists of six antigenically distinct species, including Zaire ebolavirus (represented by Ebola virus [EBOV]), Sudan ebolavirus (Sudan virus [SUDV]), Bundibugyo ebolavirus (Bundibugyo virus [BDBV]), Taï Forest ebolavirus (Taï Forest virus [TAFV]), Reston ebolavirus (Reston virus [RESV]) (Feldmann et al., 2020; Kuhn et al., 2019), and Bombali ebolavirus (Bombali virus [BOMV])(Goldstein et al., 2019). Three ebolaviruses – EBOV, BDBV, and SUDV, are responsible for severe disease and occasional deadly outbreaks in Africa posing a significant health threat. A total of 19 confirmed ebolavirus disease (EVD) outbreaks caused by EBOV have occurred, with >30,000 people infected to date and an average reported mortality rate of *∼*70%. In 2021, there are ongoing EVD outbreaks in the Democratic Republic of the Congo (DRC) and Guinea (WHO, 2021). BDBV has caused two confirmed outbreaks and infected 206 people (*∼*32% mortality rate), and SUDV has been responsible for eight confirmed outbreaks and infected 779 people (*∼*53% mortality rate) (WHO, 2021). The largest EVD epidemic to date occurred in 2013-2016 in West Africa with a total of 28,610 disease cases and 11,308 deaths reported (WHO, 2021), highlighting the urgent need for development of medical countermeasures. Monoclonal antibody (mAb) therapies have demonstrated safety and significant survival benefit in the treatment of acute EVD caused by EBOV in randomized controlled human trials (Gaudinski et al., 2019; Levine, 2019; Mulangu et al., 2019; Sivapalasingam et al., 2018), and several investigational human mAb treatments have been shown to reverse the advanced EVD in non-human primates caused by EBOV (Bornholdt et al., 2019; Corti et al., 2016; Gilchuk et al., 2020b; Pascal et al., 2018; Qiu et al., 2014), BDBV (Bornholdt et al., 2019; Gilchuk et al., 2018b), or SUDV (Bornholdt et al., 2019; Herbert et al., 2020). By 2020, two mAb-based therapeutics – ansuvimab-zykl (Ebanga) and atoltivimab + maftivimab + odesivimab-ebgn (Inmazeb) – have been developed and approved by the Food and Drug Administration (FDA) for clinical use (FDA, 2020a, b). Both of these approved antibody treatments are monospecific to EBOV, and therefore, not indicated for treatment of BDBV or SUDV infection. Identification of mAbs that cross-neutralize EBOV, BDBV, and SUDV with high potency is challenging due to the relatively high antigenic variability between these viruses (King et al., 2019). The efficacy of previously reported investigational antibody therapeutics typically is limited to only one of the three medically important ebolavirus species. The nature of future ebolavirus outbreaks cannot be predicted, however, and in a scenario of global spread viruses can mutate rapidly making available antibody treatments vulnerable to escape, as has been recently shown for SARS-CoV-2 (Starr et al., 2021; Wang et al., 2021). One approach could be to develop separate therapeutic antibody products for BDBV or SUDV or future escape variants of EBOV. It is desirable from a practical standpoint, however, to identify a single broad therapeutic spectrum antibody treatment of an equivalent or higher potency to existing monospecific antibody treatments that could be used for treatment of EBOV, BDBV, or SUDV. Therefore, ongoing efforts are needed to increase the therapeutic breadth of antibody therapies while maintaining or improving efficacy.

The ebolavirus envelope contains a single surface protein, the glycoprotein (GP), which is the key target for neutralizing mAbs (King et al., 2018; Lee et al., 2008; Lee and Saphire, 2009; Misasi and Sullivan, 2021). We previously described isolation of two broadly neutralizing human antibodies designated EBOV-515 and EBOV-442 using a human B cell hybridoma approach (Gilchuk et al., 2018a). Each of these two mAbs exhibited favorable immunological profiles, which included (1) broad reactivity for binding to GP of diverse species (EBOV, BDBV, and SUDV), (2) broad neutralization of authentic ebolaviruses (EBOV, BDBV, and SUDV), (3) recognition of distinct, non-overlapping epitopes in the GP (EBOV-515 is base-specific, and EBOV-442 is glycan-cap-specific), and (4) a high level of therapeutic protection against EBOV in mice. The antibodies EBOV-515 or EBOV-442 are analogous to the broadly-reactive mAb EBOV-520 (GP-base-specific mAb) or mAb EBOV-548 (glycan-cap-specific), respectively. We recently described a beneficial feature of the combination of EBOV-520 + EBOV-548 by showing that these two antibodies synergized for virus neutralization when combined in a cocktail and conferred therapeutic protection in EBOV-challenged rhesus macaques (Gilchuk et al., 2020b). We did not previously characterize the similar combination of EBOV-515 + EBOV-442. However, our previous studies revealed that the individual antibodies EBOV-515 and EBOV-442 had higher potency to neutralize the most antigenically distinct of the three viruses (SUDV) when compared, respectively, to the potency of EBOV-520 or EBOV-548. Also, EBOV-515 monotherapy in mice demonstrated a high level of therapeutic protection against SUDV (Gilchuk et al., 2018a; Gilchuk et al., 2020b). Given the higher potency of rEBOV-515 and rEBOV-442 against SUDV, we suggested their combination as a candidate for a “next-generation” broad therapeutic antibody cocktail. In this study we described pre-clinical development of the EBOV-515 + EBOV-442 antibody cocktail and defined the molecular basis for its pan-ebolavirus activity and efficacy.

## Results

### Activities of pan-ebolavirus candidate cocktail containing recombinant antibodies rEBOV-442 and rEBOV-515

Antibody variable gene sequences for mAbs EBOV-515 and EBOV-442 were determined, and synthetic DNAs encoding the mAbs were used to produce recombinant IgG1 in transiently-transfected Chinese hamster ovary (CHO) cells. Recombinant (r) mAbs designated as rEBOV-515 and rEBOV-442 potently neutralized EBOV, BDBV, and SUDV (**Figure 1A**) with a note that rEBOV-442 was more active against Boniface variant of SUDV (see below). To assess the Fc-mediated effector function of these antibodies to mediate killing of antigen-expressing cells, we used a stably-transfected EBOV GP-expressing SNAP-tagged 293F cell line as a target, with heterologous human PBMCs as source of effector cells and a previously described assay (Domi et al., 2018). The glycan-cap-specific antibody rEBOV-442 IgG1 exhibited a high level of cytotoxic activity, and the GP-base-specific antibody rEBOV-515 exhibited a moderate level of cytotoxic activity, relative to the control Fc-function-disabled IgG1-LALA-PG variant of rEBOV-515 (designated as rEBOV-515 LALA-PG) or an irrelevant antibody rDENV 2D22 (IgG1 isotype) specific to the dengue virus envelope (E) protein (Fibriansah et al., 2015) (**Figure 1B**). Neutralizing activity of the wild-type IgG1 and IgG1-LALA-PG variants of rEBOV-515 was equivalent (**Figure S1A**). These results taken together revealed two complementary activities exhibited by this broad antibody cocktail, specifically the Fc-mediated effector functions mediated by rEBOV-442 and the potent neutralizing activities mediated by rEBOV-442 and rEBOV-515 (**Figure 1C**).

**Figure 1.**
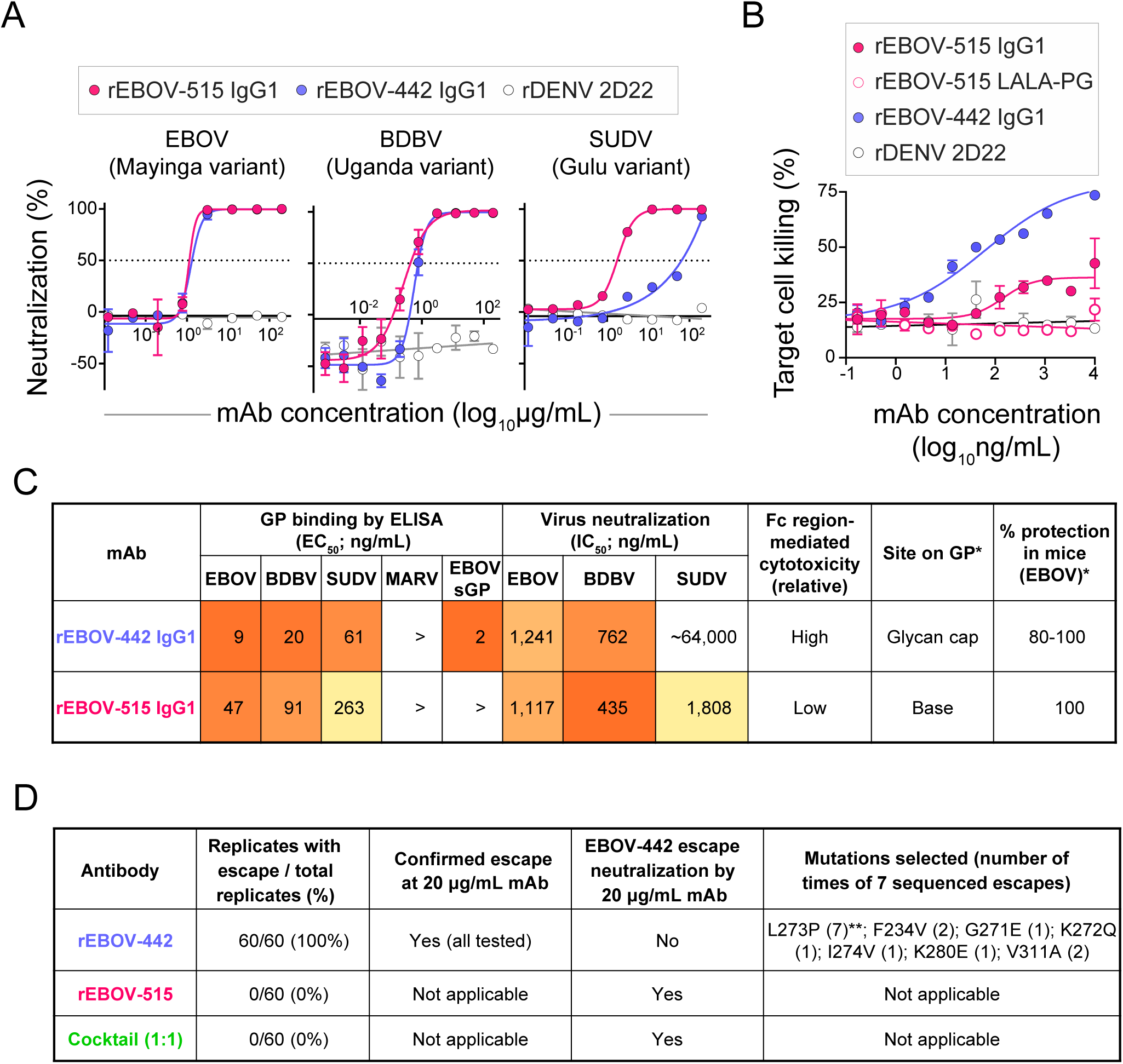
Functional activities of pan-ebolavirus cocktail candidate antibodies rEBOV-442 and rEBOV-515. (A) EBOV, BDBV, or SUDV neutralization. Biosafety level 4 recombinant ebolaviruses encoding enhanced green fluorescent protein (eGFP) were incubated with increasing concentrations of recombinantly produced purified mAbs, and infection was determined at 3 days after inoculation by measuring eGFP fluorescence in cells. Mean ± SD of technical triplicates from one experiment are shown. (B) *In vitro* killing capacity mediated by the Fc regions of IgG1-engineered variants of mAbs measured by rapid fluorometric antibody-mediated cytotoxicity (RFADCC) assay. Human PBMCs (effector cells) were incubated with a SNAP-tagged EBOV GP-expressing 293F cell line as a target in the presence of increasing concentrations of purified recombinant mAbs, and cytotoxic activity was measured by flow cytometry. Dengue virus human antibody rDENV 2D22 served as a negative control. The dotted line indicates assay background. Mean ± SD of technical duplicates from one experiment are shown. (C) Heat map summarizing reactivity breadth and potency of rEBOV-442 and rEBOV-515. * indicates data determined in our previous work using hybridoma-cell-secreted mAbs (Gilchuk et al., 2018). (D) Results of viral selections with individual antibodies or the cocktail, indicating the number of replicates with escape out of total number tested, resistance of selected escape variants to individual antibodies or the cocktail, and the selected mutations in the GP that escape neutralization by rEBOV-442. The escape selection was performed using infectious chimeric VSV expressing EBOV (Mayinga variant) GP. ** indicates the prevalent mutation L273P that was identified in all sequenced escape viruses. See also **Figure S1**.

One benefit of using a cocktail of two or more neutralizing antibodies that bind to non-overlapping regions of the viral antigen is to reduce the risk of viral escape from neutralization that is inherent in monotherapy approaches (Baum et al., 2020; Misasi and Sullivan, 2021; Yewdell et al., 1979). To demonstrate this feature for this cocktail directly, we next used a recombinant infectious vesicular stomatitis virus (VSV) expressing EBOV GP in place of the endogenous VSV glycoprotein (VSV/EBOV GP) to select for GP mutations that escape antibody neutralization. We assessed rEBOV-442, rEBOV-515, or 1:1 mixture of both antibodies (half maximal inhibitory concentration [IC_50_] <1.5 μg/mL against EBOV for individual mAbs and the cocktail) and used a high-throughput quantitative real-time cellular analysis assay (RTCA) to select viral variants that can escape neutralization (Gilchuk et al., 2020b; Greaney et al., 2021). We selected for escape at a single saturating antibody concentration of 20 μg/mL using *∼*20,000 plaque forming units of VSV/EBOV GP per well (*∼*1 multiplicity of infection [MOI]) and performing 60 replicates for each sample (**Figure 1D**). For rEBOV-442, this process selected viral variants that we confirmed escaped neutralization at 20 μg/mL of rEBOV-442 but were neutralized by rEBOV-515 or a mixture of rEBOV-442 and rEBOV-515 (**Figure 1D, S2B**). We sequenced the viral gene encoding the GP of rEBOV-442-selected escape viruses and identified seven distinct point mutations, including the previously identified escape mutation L273P (Gilchuk et al., 2018a) that appeared in all sequenced escape viruses in this study (**Figure 1D**). The escape mutation L273P likely pre-existed in the virus stock at a low frequency explaining the observed rate of escape from rEBOV-442 neutralization. In all cases, escape viruses carried GP mutations in the binding site of rEBOV-442 (**Figure S1C**). For rEBOV-515 or the cocktail, escape variants were not detected in any of the 60 replicate experiments (**Figure 1D**). Together these results demonstrated a high resistance to viral escape for the cocktail of the two broad antibodies rEBOV-442 and rEBOV-515, which acts by both neutralizing and Fc-mediated activities.

### Differential requirements for the Fc regions of antibodies rEBOV-442 and rEBOV-515 for therapeutic protection in mice

To define the contribution of Fc-mediated effector functions to protection *in vivo*, each mAb of the candidate therapeutic cocktail was assessed as an IgG1 protein (the original subclass of antibody secreted by the hybridoma cell line and a functionally competent form of mAb) and as IgG1 LALA-PG protein (which is disabled for Fc function and fully silent in mice). We challenged groups of mice with mouse-adapted EBOV (EBOV-MA) on day 0 and administered antibodies 1 day later. Previously we have shown that for low-dose treatment (1 mg/kg), the Fc-disabled LALA variant of the GP base-specific mAb EBOV-520 offered a higher level of therapeutic protection in mice against EBOV (60% survival) when compared to that mediated by the wild-type (*wt*) IgG1 EBOV-520 (0% survival; (Kuzmina et al., 2018)). In a new study conducted here, the rEBOV-515 LALA-PG antibody variant offered complete protection (100% survival) against EBOV challenge in mice at a 1 mg/kg therapeutic dose, while the *wt* IgG1 variant showed partial protection (80% survival) (**Figure 2**). This finding demonstrated a higher *in vivo* potency of the broad GP base-specific mAb rEBOV-515 when compared to the previously reported effect of the broad GP base-specific mAb rEBOV-520, and suggested that rEBOV-515 LALA-PG is a preferable Fc variant for the cocktail of two Fc variants tested. Similarly, we compared the *in vivo* potency of *wt* IgG1 rEBOV-442 and rEBOV-442 LALA-PG, which were administered at a 5 mg/kg dose one day after EBOV challenge (**Figure 2**). Cytotoxicity assay results (**Figure 1B**) suggested that the glycan cap antibody rEBOV-442, in addition to possessing neutralizing activity, also exhibited substantial Fc effector function-mediated activity. In agreement with this finding, the results of our *in vivo* study showed a transient (2-4 days post-infection) improvement in weight (*i.e.*, reduced loss) with *wt* rEBOV-442 IgG1, which was associated with improved protection (100% survival in the *wt* antibody-treated group when compared to 80% in the rEBOV-442 LALA-PG-treated group) (**Figure 2**). Together, these data revealed differing requirements for the Fc regions of rEBOV-442 and rEBOV-515, demonstrated a high *in vivo* potency of the antibodies tested, and justified the inclusion of rEBOV-515 as a LALA-PG variant and rEBOV-442 as a *wt* IgG1 variant in a new broad and protective ebolavirus two-antibody cocktail.

**Figure 2.**
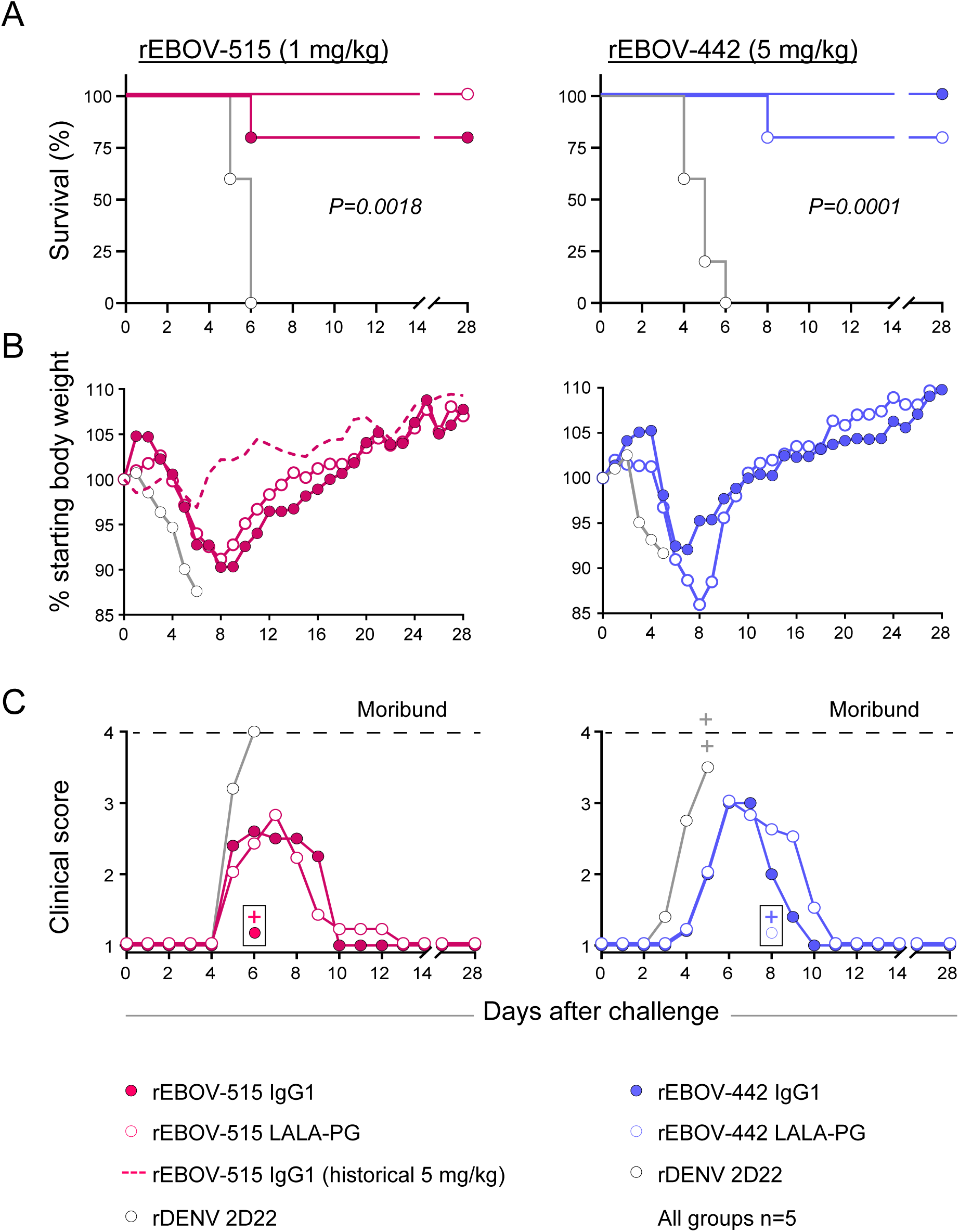
Differential requirements for the Fc regions of antibodies rEBOV-442 and rEBOV-515 for optimal therapeutic protection in mice. C57BL/6 mice were challenged with mouse-adapted EBOV-MA, treated at 1 dpi with wild-type IgG1 or the Fc-effector function disabled IgG1 LALA-PG variant of antibodies rEBOV-442 (5 mg/kg) and rEBOV-515 (1 mg/kg), and monitored for 28 days. (A) Kaplan-Meier survival plot. The overall difference in survival between the groups was estimated using two-sided log-rank (Mantel–Cox) test. (B) Weight change. (C) Clinical score. +, animal found dead prior to reaching the pre-determined clinical score. Mean values are shown in (B-C), and data represent one experiment with five mice per group. Dengue virus-specific antibody rDENV 2D22 was used as a control. A historical control for protection from weight loss with 5 mg/kg of wild-type rEBOV-515 treatment (dotted line in B) is shown for comparative purposes.

### Synergistic activity of antibodies rEBOV-442 and rEBOV-515 in the cocktail

Antibodies in a cocktail directed to a common viral protein may recognize antigen in a synergistic, additive, or antagonistic manner. We first characterized the effect mediated by the mixture of two antibodies on ebolavirus GP binding. Serially-diluted Alexa-Fluor-647-labeled antibody rEBOV-442 IgG1 or rEBOV-515 LALA-PG was titrated into serially-diluted unlabeled partner antibody to generate two pairwise combinatorial matrices of two antibodies in the mixture (**Figure S2A, B**). Binding of the Alexa-Fluor-647-labeled antibody from each combination was assessed by flow cytometric analysis using a Jurkat cell line that was stably transduced to display EBOV GP on the surface as described before (Gilchuk et al., 2020b). The values were calculated as the percent of the relative fluorescence signal caused by the highest concentration of respective fluorescently-labeled antibody alone (**Figure S2A, B**), and then compared with the expected responses calculated using the ZIP synergy-scoring model (Ianevski et al., 2020). The synergy score can be interpreted as the average excess response (percent change) for an antibody combination (Ianevski et al., 2020). The comparison revealed that the combination of antibodies rEBOV-442 and rEBOV-515 was synergistic, and each antibody reciprocally enhanced binding of the partner antibody in the mixture (**Figure S2B, C**). This finding contrasted with the unidirectional GP binding synergy observed in the cocktail of the other broad antibodies rEBOV-520 and rEBOV-548 (Gilchuk et al., 2020). Binding enhancement increased steadily with increasing of antibody concentration, indicating more contribution from synergy at higher concentrations of individual Abs in the cocktail (**Figure S2B, C**).

We next characterized the effect by the mixture of two antibodies on ebolavirus neutralization. Using a recently developed real-time cell analysis (RTCA) cellular impedance assay that quantifies virus-induced cytopathic effects and infectious chimeric VSV/ebolavirus GP viruses (Gilchuk et al., 2020a; Gilchuk et al., 2020b), we showed efficient and dose-dependent neutralization of VSV/EBOV-GP, /BDBV-GP, or /SUDV-GP viruses by the 1:1 antibody mixture (**Figure 3A**). Then, we titrated serially-diluted rEBOV-442 into serially-diluted rEBOV-515 to generate a pairwise combinatorial matrix of two antibodies in the mixture and assessed each combination for neutralization of VSV/EBOV-GP, /BDBV-GP, or /SUDV-GP viruses (**Figure 3B**). Data were analyzed using the ZIP synergy scoring model. This analysis identified areas in the neutralization matrix (showing mAb ratios) with the most profound combination effect and suggested broad synergistic activity in the cocktail for the mixture of rEBOV-442 and rEBOV-515 (**Figure 3C**).

**Figure 3.**
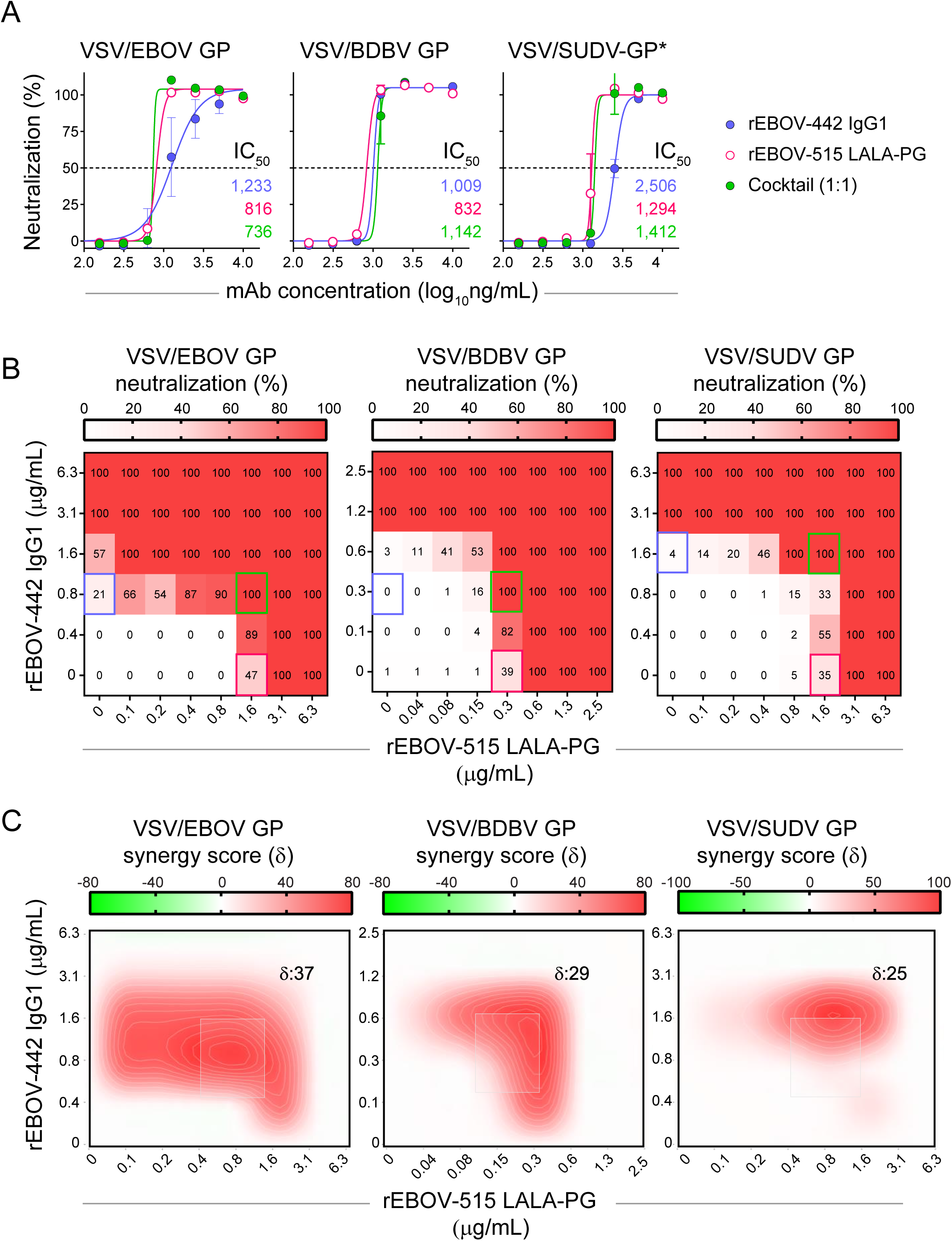
Broad and synergistic neutralizing activity mediated by the cocktail of rEBOV-442 and rEBOV-515. Neutralizing activity of individual antibodies or their mixture was assessed using infectious chimeric VSV expressing EBOV (Mayinga variant), BDBV (Uganda variant), or SUDV (Boniface variant) GP and real-time cell analysis (RTCA) assay. (A) Neutralization of VSV/EBOV GP, VSV/BDBV GP, or VSV/SUDV GP by rEBOV-442 alone, rEBOV-515 alone, or a 1:1 mixture of rEBOV-442 and rEBOV-515. (B) Serially-diluted rEBOV-442 was titrated into serially-diluted rEBOV-515 to generate a pairwise combinatorial matrix of two antibodies in the mixture. The matrix shows neutralization dose-response data for VSV/EBOV GP, VSV/BDBV GP, or VSV/SUDV GP, by indicated concentrations of rEBOV-442 and rEBOV-515. Axes denote the concentration of each antibody, with the percent neutralization shown in each square. The heat map denotes a gradient of 0 (white) to 100% (red) neutralization. Examples of neutralization by rEBOV-515 alone (raspberry box) or rEBOV-442 alone (blue box) in comparison to a combined indicated concentration of two mAbs in the cocktail (green box) are shown. (C) Synergy distribution map generated from the dose-response neutralization matrix in (B). Red color indicates areas in which synergistic neutralization was observed; shaded grey box indicates the area of maximum synergy between the two monoclonal antibodies, and the *δ*-score for this area is shown. The *δ*-score is a synergy score: values < -10 indicate antagonism; values -10 to 10 indicate an additive effect; values >10 indicate synergy. Data in (A-C) are from a representative experiment performed in technical duplicate and repeated twice. See also **Figure S2**.

### Protective pan-ebolavirus combination therapy of nonhuman primates by antibodies rEBOV-442 and rEBOV-515

Next, we tested the efficacy of the cocktail of rEBOV-515 LALA-PG + rEBOV-442 IgG1 in nonhuman primate (NHP) challenge models for each of the three ebolaviruses, EBOV (Kikwit variant), BDBV (Uganda variant), and SUDV (Gulu variant). We used rhesus macaque EBOV and SUDV lethal challenge models and a cynomolgus monkey BDBV lethal challenge model, which recapitulate many key features of EVD in humans (Bennett et al., 2017; Geisbert et al., 2015). Animals were assigned to three treatment groups of five animals per group. After intramuscular challenge with a lethal target dose of 1,000 plaque-forming units (PFU) of EBOV, BDBV, or SUDV, all NHPs of the treatment group received intravenously two 30 mg/kg doses of the cocktail (a 2:1 mixture of rEBOV-515 LALA-PG and rEBOV-442 IgG1) spaced 3 days apart (given days 3 and 6 after EBOV, days 6 and 9 after BDBV, or days 4 and 7 after SUDV inoculation). We chose a 2:1 antibody ratio in the cocktail based on the high level of synergy identified for this antibody ratio from the *in vitro* synergy distribution maps (**Figure 3; S2**) and due to higher neutralizing potency of rEBOV-515 against SUDV when compared to the potency of rEBOV-442 (**Figure 1A**). For each challenge cohort, an additional animal was studied as a contemporaneous control and was left untreated. The untreated control animals developed a high clinical score and succumbed to disease on day 6, 14, or 10 after viral challenge with EBOV, BDBV, or SUDV, respectively. The two-dose therapeutic cocktail treatment provided complete pan-ebolavirus protection of NHPs from mortality and clinical signs of EVD (**Figure 4A, B**). We next assessed changes in blood chemistries and blood cell composition that are typically associated with EVD to further characterize the efficacy of the mAb cocktail treatment (**Tables S1-6**). The liver enzymes alkaline phosphatase (ALP) and gamma glutamyl transferase (GGT), which are indicators of EVD (Qiu et al., 2014), were elevated in untreated NHPs at the peak of the disease (**Figure 4B, C**). Treated animals did not show signs of acute liver injury after the treatment, displaying low amounts of ALP, GGT, creatinine (CRE), and the other blood chemistries when compared with those of untreated NHP (**Figure 4C; Tables S4-6**).

**Figure 4.**
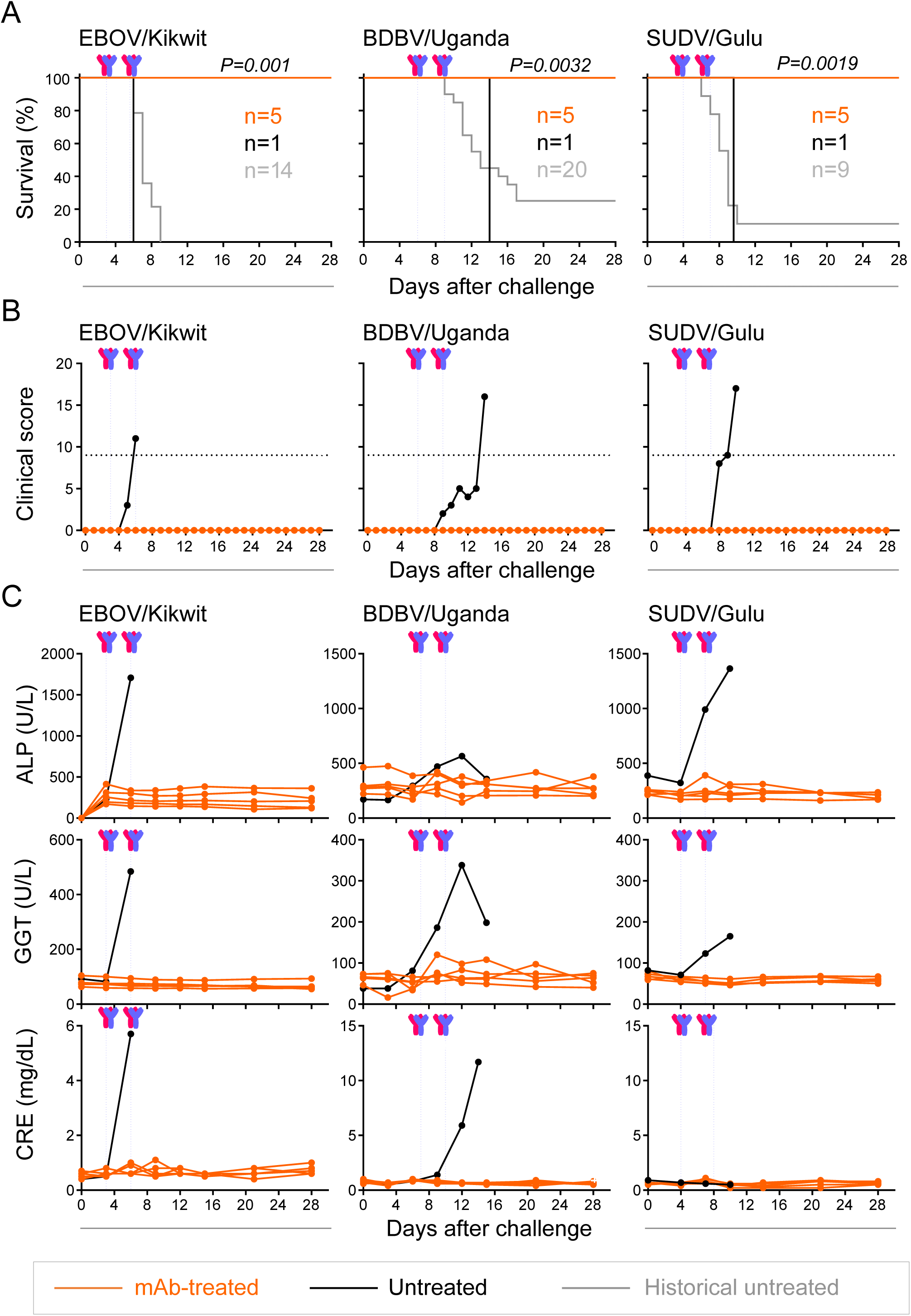
The cocktail treatment provides pan-ebolavirus protection of nonhuman primates against disease. Rhesus macaques were inoculated with a lethal dose of the EBOV Kikwit or SUDV/Gulu viruses intramuscularly (i.m.) on day 0 and were treated with total 30 mg/kg of the cocktail (1:2 mixture of rEBOV-442 and rEBOV-515) intravenously on 3 and 6 dpi (EBOV/Kikwit; n = 5 per cohort), or 4 and 7 dpi (SUDV/Gulu; n = 5 per cohort). Cynomolgus monkeys were inoculated with a lethal dose of the BDBV/Uganda i.m. on day 0 and were treated with a total dose of 30 mg/kg of the cocktail (1:2 mixture of rEBOV-442 and rEBOV-515) intravenously on 6 and 9 dpi (n = 5 per cohort). The contemporaneous control was an untreated NHP challenged with the virus (n = 1 for each cohort). One experiment was performed. (A) Kaplan-Meier survival plot. The historical untreated controls (grey) are shown for comparative purposes (see Methods). The proportion surviving at day 28 after viral challenge in the treated cohort was compared to the respective historical untreated cohort using a 2-sided exact unconditional test of homogeneity. (B) Clinical score. (C) Selected blood chemistry measurements: ALP, alkaline phosphatase; GGT, gamma-glutamyl transpeptidase; CRE, creatinine. Antibody treatment times are indicated with blue dotted vertical lines. Orange curves indicate treated, and black indicate untreated animals in (A) to (C). The black dotted line in (B) indicates the clinical score threshold for euthanasia. See also **Tables S1-6**.

At the time of first treatment with the cocktail (day 3 after EBOV, day 6 after BDBV, or day 7 after SUDV inoculation), 14 NHPs from the treatment-designated group (all except one animal in EBOV-challenged group) and the control untreated NHPs were viremic, with virus titers that ranged from 5.1 to 10.6 log_10_ genome equivalents (GEQs) per mL of plasma, as measured by quantitative reverse-transcription PCR (qRT-PCR) (**Figure 5A**). A plaque assay that detects infectious virus revealed viremia in all 18 animals, with viral titers ranging from 0.7 to 7.4 log_10_ plaque forming units (PFU) per mL of plasma at the time of first treatment with the cocktail (**Figure 5B**). This finding confirmed active ebolavirus infection in all NHPs before treatment. At the time of the second antibody cocktail treatment (day 6 after EBOV, days 9 after BDBV, or day 7 after SUDV inoculation), each untreated control NHP from a respective EBOV, BDBV, or SUDV challenge cohort developed high viremia >8 log_10_ GEQ/mL and >5 log_10_ PFU/mL (**Figure 5 A, B**). Concordant with full protection from disease and death, none of the 15 treated NHPs had detectable infectious virus in the plasma at that time (limit of detection [LOD] = 5 PFU/mL), and viral genomes were not detected in 12 of those NHPs 3.7 log_10_ GEQ/mL at that time (**Figures 4, 5**). The three remaining NHPs that were from the BDBV inoculation cohort and that showed slightly delayed clearance of viral genomes in plasma no longer had detectable virus by day 15 after viral challenge. Analysis of various tissues harvested at study endpoint from treated animals (day 28 after inoculation with EBOV and SUDV or day 35 after inoculation with BDBV) and untreated animals (day 6, 14, or 10 after inoculation with EBOV, BDBV, or SUDV, respectively) confirmed virologic protection mediated by the cocktail treatment (**Figure S3**).

**Figure 5.**
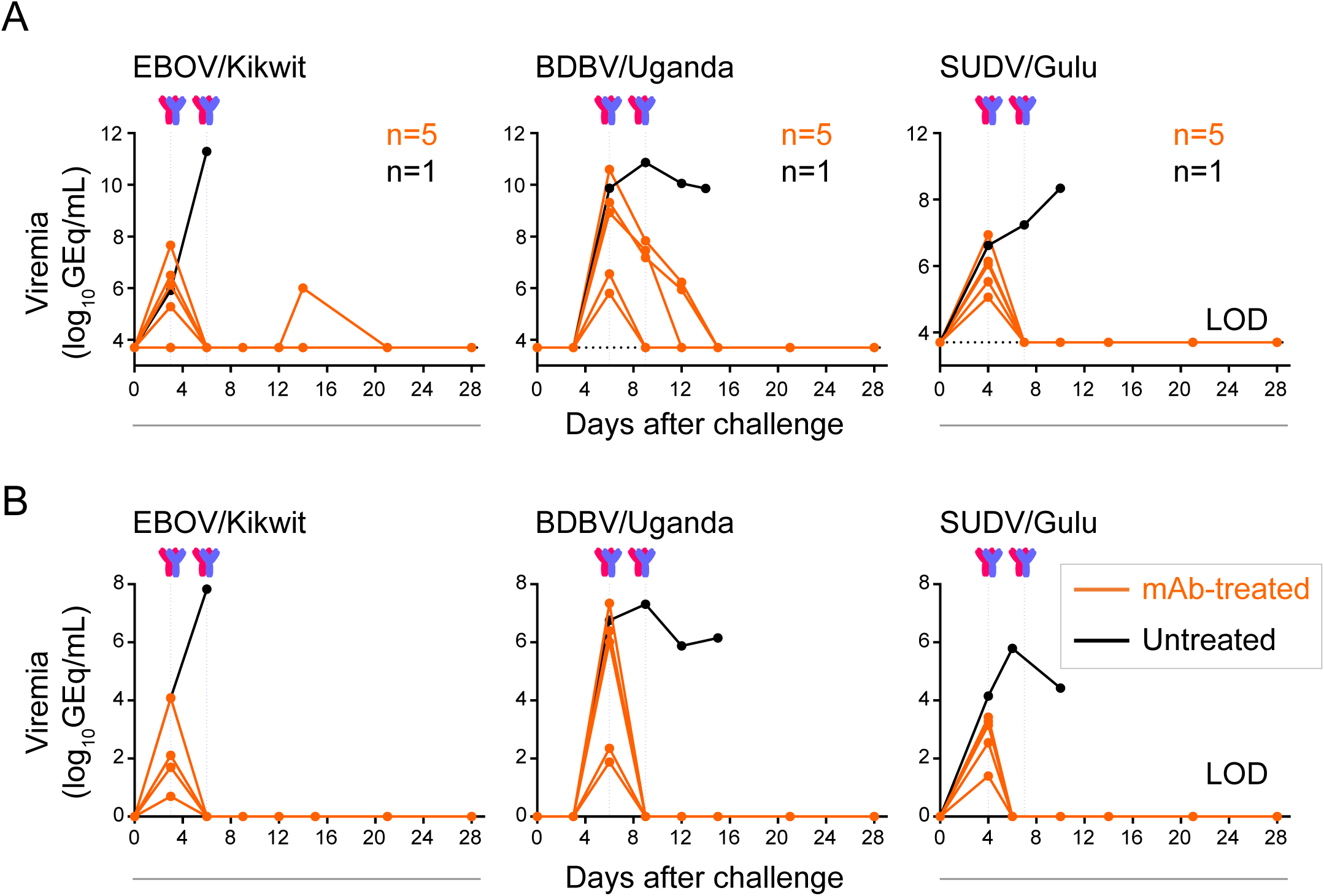
The cocktail treatment provides pan-ebolavirus protection of nonhuman primates against viremia. Blood viral loads were assessed from individual animals that were challenged with EBOV/Kikwit, BDBV/Uganda, or SUDV/Gulu and treated with the two-antibody cocktail as described in Figure 4. (A) Kinetics of blood viral load determined for genome equivalents (GEq) using qRT-PCR. (B) Kinetics of infectious virus blood viral load as determined by plaque assay. Orange curves indicate treated, and black indicate untreated animals. Antibody treatment times are indicated with blue dotted vertical lines. The black dotted line indicates the limit of detection (LOD), which was 3.7 log_10_GEq/mL (A) or 5 PFU/mL (B). Each measurement in (A-B) represents the mean of technical duplicates. See also **Figure S3**.

Together these results showed a high therapeutic efficacy of the cocktail of rEBOV-515 LALA-PG + rEBOV-442 IgG1 to treat and revert disease caused by primary ebolaviruses that are responsible for outbreaks in humans – EBOV, BDBV, and SUDV.

### Structural basis for the efficacy and broad ebolavirus neutralization by the cocktail

We previously reported the molecular determinants of the GP binding for several glycan cap-specific antibodies, including the broad antibody rEBOV-442 (Murin et al., 2021), and defined the structural basis of synergy for the pair of broad antibodies rEBOV-520 and rEBOV-548 (Gilchuk et al., 2020b). In this new study, we focused on studies of the determinants of rEBOV-515 binding to elucidate the structural basis for the neutralization breadth and efficacy mediated by pan-ebolavirus cocktail of rEBOV-515 + rEBOV-442. We generated a complex of both rEBOV-515 and rEBOV-442 Fab with mucin-deleted EBOV GP from the Makona variant (EBOV GPΔMuc/Mak) and solved structures by cryogenic electron microscopy (cryo-EM) (**Table S7;** **Figure 6A** **and S4**).

**Figure 6.**
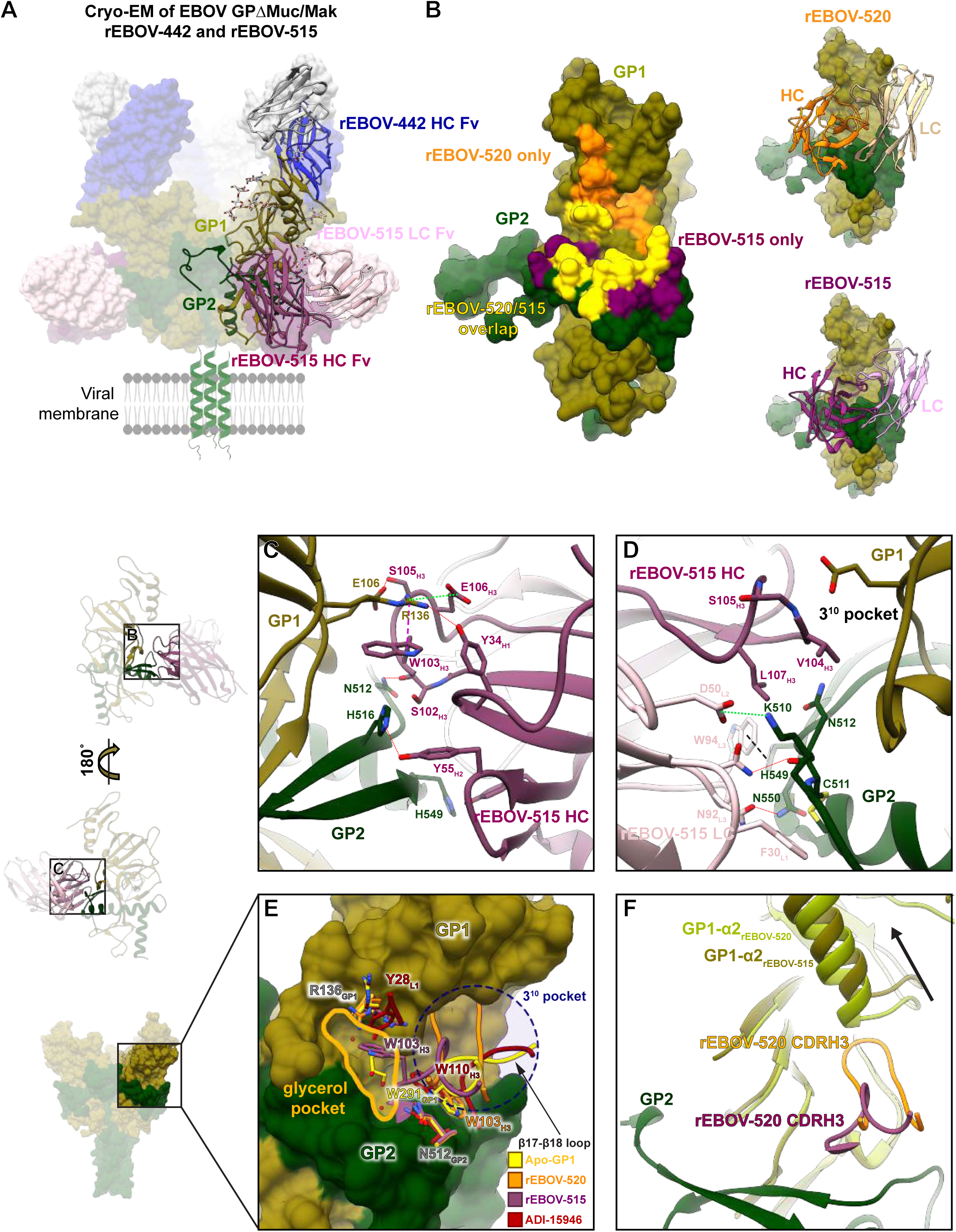
rEBOV-515 binds to the major site of vulnerability in the GP base region in a distinct manner. (A) Cryo-EM structure of rEBOV-442 (heavy chain in blue and light chain in grey) and rEBOV-515 (heavy chain in maroon and light chain in pink) Fab bound to EBOV GPΔMuc/Mak. A side view in relation to the viral membrane is shown. Fab constant domains were excluded by masking. (B) The predicted contact surface of broadly neutralizing antibodies rEBOV-515 and rEBOV-520 on the surface representation of the EBOV GPΔMuc/Mak monomer model (PDB: 5JQ3). Non-overlapping contact surfaces for rEBOV-515 or rEBOV-520 are shown, respectively, in maroon or orange, and overlapping contact surface of both antibodies is shown in yellow. (C) Contact residue details of the rEBOV-515 heavy chain Fab interactions with the base of GP. CDRH3 contacts include a backbone-mediated hydrogen bond at S102_H3_ to N512_GP2_, a key contact that links the β17-β18 loop to the base of the IFL via W291 in unliganded GP1. Contacts near the 3^10^ pocket include a potential hydrogen bond between S105_H3_ and E106_GP1_ that forms when a large portion of the CDRH3 loop displaces the β17-β18 loop. Within the CDRH2 loop, Y52_H2_ and Y55_H2_ make potential hydrogen bonds via their hydroxyl groups to H549_GP2_ or H516_GP2_ on GP2, respectively. One of the most extensive rEBOV-515 contacts is from W103_H3_, which forms a strong cation-pi bond with R136_GP1_, allowing R136 to make additional hydrogen bonds with Y34_H1_ and a salt bridge with E106_H3_. (D) Epitope details of the rEBOV-515 light chain interactions with the base of GP. The rEBOV-515 light chain makes contacts with all three CDRs exclusively within GP2. Contact features include a potential hydrogen bond between N32_L1_ and the backbone of C511_GP2_, and a salt bridge between D50_L2_ and K510_GP2_. Pi cation interactions at W94_L3_ with H549_GP2_ (black dashed line) and an additional potential hydrogen bond at N92_L3_ with N550_GP2_ provide additional stabilizing interactions. (E) Comparison of antibody CDRH3 loops that bind in and around the 3^10^ pocket (blue dashed circle) and putative glycerol pocket (solid yellow line). rEBOV-520 and ADI-15946 replace and mimic residue W291_GP1_ (that anchors down the β17-β18 loop in apo-GP) with a tryptophan from their CDRH3 loops. rEBOV-515 uses an analogous tryptophan residue (W103_H3_) to contact R136_GP1_ via a strong cation-pi interactions bond, which causes a shift in the rotamer of R136_GP1_. W103_H3_ from rEBOV-515 also accesses a pocket that is occupied by a glycerol cryoprotectant molecule in the unliganded crystal structure of GP, which is also occupied by Y28_L1_ from ADI-15946. (F) A shift in the placement of the GP1 α2 helix that cased by the CDRH3 from rEBOV-520 but is lacking in rEBOV-515 binding due to a shorter CDR loop is shown. Red dotted lines: hydrogen bonds; green dashed lines: salt bridge; purple dashed line: cation-pi interaction; black dashed line: carbon-pi or aromatic interactions. rEBOV-520-GP PDB: 6UYE; ADI-15946-GP PDB: 6MAM. See also **Figures S4-6 and Table S7**.

The interface between GP and rEBOV-515 in this structure was resolved to ∼3 to 3.5 Å resolution (**Figure S4**), allowing for confident modeling of most of the side-chain residues. rEBOV-515 binds to the base of the internal fusion loop (IFL), directly below the glycan cap in GP1 (**Figure 6A**). There is significant overlap of the rEBOV-515 epitope with that of rEBOV-520, but rEBOV-515 has more extensive contacts in GP2 and less in GP1 (**Figure 6B**). Most EBOV-515 contacts are mediated by the heavy chain (HC), with all three complementarity determining regions (CDRs) contributing to binding the base of the IFL as well as a portion of GP1 near the 3^10^ pocket (**Figure 6C; S5A-B, S6**). The CDRH3 contributes the most extensive contacts, including those that displace the descending β17-β18 loop of the glycan cap, making contacts near the 3^10^ pocket, as well as a strong cation-pi bond with R136_GP1_ (**Figure 6C**). The rEBOV-515 light chain (LC) also makes contacts with all three CDRs exclusively within GP2 (**Figure 6D**; **S5A**). Together, this analysis showed that rEBOV-515 forms a strong interaction of complementary hydrophobic (**Figure S5B**) and electrostatic (**Figure S5D**) surfaces for binding to this epitope within GP.

Neutralizing potency may depend on the ability of antibodies to remain bound to viral GP within the acidic environment of late endosomes, which is where pH-dependent cleavage occurs to expose the Niemann-Pick C1 (NPC1) receptor binding site (Carette et al., 2011; Chandran et al., 2005). Given that rEBOV-515 strongly interacts with the EBOV GP, we assessed binding of this antibody at varying pH to recombinant EBOV, BDBV or SUDV GPs by ELISA. We used the broadly reactive base-specific antibody rEBOV-520 for comparison. rEBOV-515 and rEBOV-520 bound equivalently to the GP of each of the three ebolaviruses at neutral pH 7.4, and rEBOV-515 remained bound at acidic pH of 5.5 or 4.5, while rEBOV-520 lost binding activity (**Figure S7A**).

The rEBOV-515 binding site partially overlaps with those of other reported broad antibodies that bind to the 3^10^ pocket, including rEBOV-520 (Gilchuk et al., 2020b) and ADI-15946 (West et al., 2019) (**Figure 6B, E**). Each of these antibodies relies upon E106 and R136 in GP1 and several residues at the base of the IFL, using long CDRH3 loops that mimic and replace the β17-β18 loop of the GP1 in apo-GP structure (**Figure 6E**). However, characterization of the rEBOV-515 binding site revealed several unique features. rEBOV-515 uses a single CDR loop to simultaneously mimic and displace the β17-β18 loop and to bind to R136 in GP1. This feature allows rEBOV-515 to access a region that is occupied by a glycerol cryoprotectant molecule in the unliganded crystal structure of GP (Protein Data Base [PDB] 5JQ3; 5JQB) that we defined as “glycerol pocket” (**Figure 6E**). In addition, this mechanism facilitates greater interaction of EBOV-515 with the cleavage loop compared to that caused by rEBOV-520, which may explain the high GP cleavage-inhibiting activity of EBOV-515 (**Figure S7B**). Further, a distinct pose of rEBOV-515 binding into the 3^10^ pocket avoids clashes with the *α*2 helix of the glycan cap (**Figure 6F**), unlike the binding of rEBOV-520 to the GP, which requires the *α*2 helix shift (Gilchuk et al., 2020b). This finding suggested that the binding site of rEBOV-515 on the intact GP molecule is more accessible than the rEBOV-520 binding site, which could explain the higher neutralizing potency of rEBOV-515 when compared to the potency of rEBOV-520 against the virus carrying the intact GP. Of note, we have shown previously that GP cleavage, which removes the glycan cap along with the β17-β18 loop, resulted in enhanced binding to cell-surface-displayed cleaved GP and increased neutralizing potency for both rEBOV-515 and rEBOV-520 (Gilchuk et al., 2018a; Gilchuk et al., 2020b).

There are notable differences in degree of pan-ebolavirus neutralization and protection by each of three reported broadly-reactive human antibodies despite recognition of partially overlapped epitopes they recognize on the GP base (**Figure 6B**). ADI-15946 did not fully neutralize most antigenically distinct SUDV (Wec et al., 2017; West et al., 2019), rEBOV-520 neutralized SUDV with modest potency but showed weak protection against SUDV in mice, and rEBOV-515 potently neutralized each of three viruses and protected against SUDV in mice (Gilchuk et al., 2018a; Gilchuk et al., 2020b). Our structural analysis allowed us to compare the epitope footprint and contacts for each of these antibodies in relation to sequence conservation between the three major ebolavirus (**Figure 7; S6**), providing insight that may help explain differences in neutralization breadth. ADI-15946 makes fewer contacts with GP1 but more contacts to the N-terminus of GP2, a region that is less conserved that may explain weak SUDV neutralization. Conversely, rEBOV-520 makes more extensive contacts with GP1, as this region is generally less conserved. rEBOV-515 relies on minimal contacts with GP1, except in regions that are completely conserved, including R136 and E106. Taken together with a more conserved footprint, these observations provide a molecular basis for the superior breadth of neutralization and protection exhibited by rEBOV-515 over similar antibodies.

**Figure 7.**
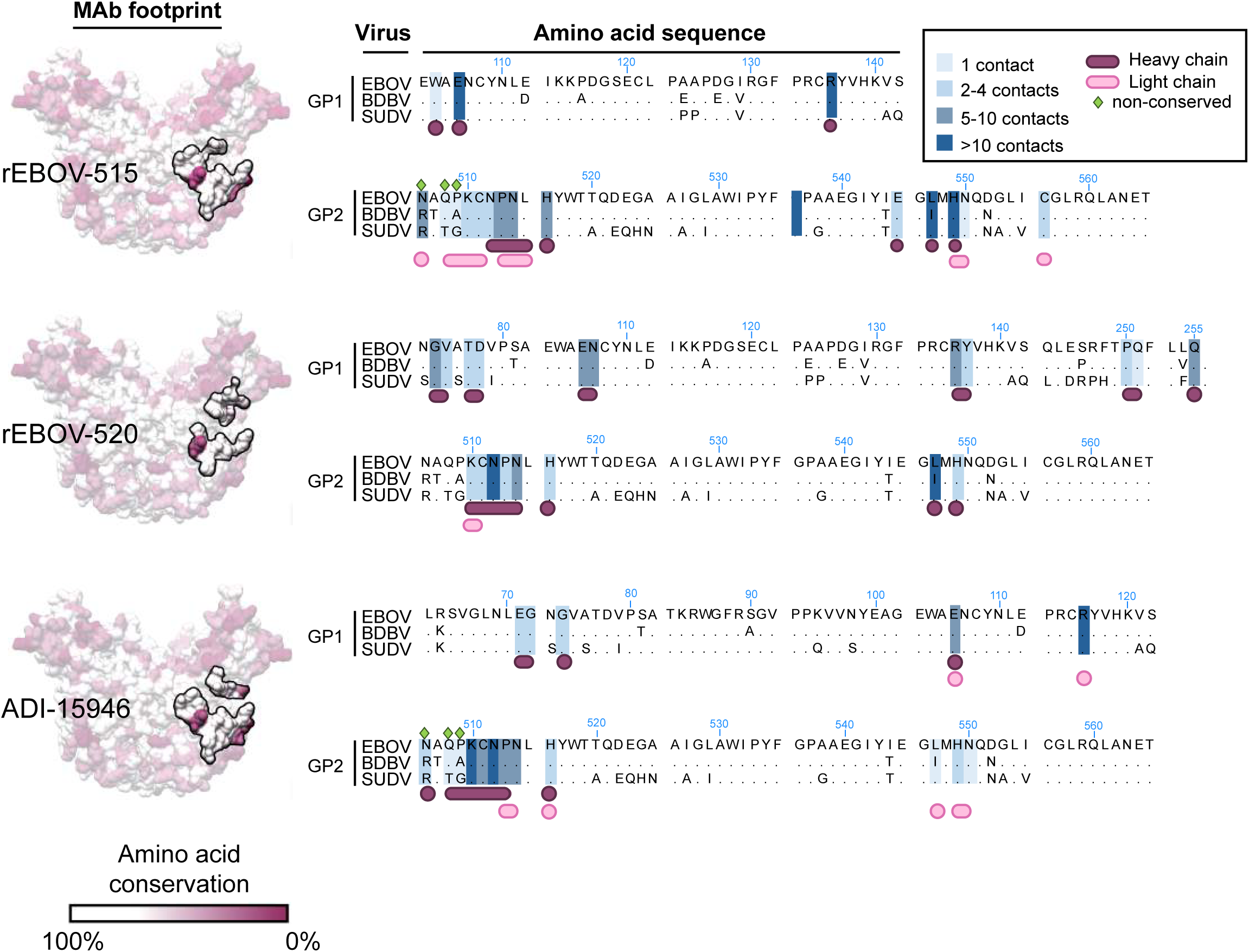
Conservation of the binding sites for broadly-reactive antibodies rEBOV-515, rEBOV-520, and ADI-15946. Apo-GP is surface-rendered according to residue conservation on the left, with no conservation in dark purple (0%) and complete conservation in white (100%). The corresponding antibody footprints are highlighted. On the right are aligned sequences of the interacting regions on GP from the three major ebolaviruses, EBOV, BDBV, and SUDV. Total contacts for each residue at 4 Å distance or less were determined (see **Table S7**) and residues are highlighted in blue according to the number of contacts, with darker blue indicating more contacts and thus a higher likelihood for contributing critically to binding. Residues that are variable are marked with a green diamond. HC contacts are indicated below in dark purple and LC contacts in pink.

## Discussion

Here we report comprehensive studies of a new pan-ebolavirus antibody combination treatment with a well-defined and complex molecular basis of broad neutralization and potency. The cocktail exhibited high therapeutic efficacy against all three medically important ebolaviruses in nonhuman primates. The discovery and recent approval of human antibody-based therapeutics represents a landmark achievement in the development of EVD medical countermeasures. The clinical trials in the DRC outbreak, however, highlighted substantial gaps remained in improving the treatment of acute EVD (Iversen et al., 2020). The greatest benefit of antibody treatment in patients was mainly in those receiving early therapy, and only moderate benefit was observed in severely ill patients (Levine, 2019). Another remaining challenge for implementation of the current regimens is the logistical complexity of intravenous drug administration in the field, which may limit widespread application of antibody therapy in future outbreak scenarios.

Given the difference in efficacy mediated by different antibody drugs that was observed for advanced EVD treatments, antibody potency likely is a key contributing determinant of treatment efficacy. Inmazeb and Ebanga demonstrated higher efficacy when compared to that of the antibody cocktail ZMapp™ in clinical trials (Levine, 2019). Inmazeb is a three-antibody cocktail based on REGN-EB3 antibody sequences (Pascal et al., 2018), Ebanga is a monotherapy based on the antibody mAb114 (Corti et al., 2016), and ZMapp™ is a cocktail of three murine-human chimeric mAbs (Qiu et al., 2014). A comparison with the historical NHPs studies with REGN-EB3 (3 x 50 mg/kg dose at d5, 8, and 11 after exposure) (Pascal et al., 2018), mAb114 (3 x 50 mg/kg dose at d1, 2, and 3 after exposure) (Corti et al., 2016), and ZMapp™ (3 x 50 mg/kg dose at d3, 6, and 9 after exposure) (Qiu et al., 2014) suggested equivalent or likely higher potency of our EBOV-442 IgG1 + EBOV-515 LALA-PG cocktail treatment (2 x 30 mg/kg at d3 and 6 after exposure) against homologous EBOV. More studies are needed to compare available treatments and to determine if increasing antibody therapeutic potency would benefit clinical outcomes in treatment of severely ill patients and/or allow for rapid and more practical treatment by an alternative (intramuscular or subcutaneous) route of antibody delivery.

Anticipation of future EVD outbreaks requires consideration of therapeutic breadth. As an added feature, the two-antibody cocktail we describe here offered an increased therapeutic breadth that extended to protection against BDBV and SUDV in NHP models. One other investigational human mAb cocktail that demonstrated broad efficacy in NHPs, MBP134^AF^, has been described (Bornholdt et al., 2019). MBP134^AF^ is comprised of antibody ADI-15878 and a derivative of antibody ADI-15946 (defined as ADI-23774), which was engineered to improve its specificity against SUDV GP (Wec et al., 2019). A comparison of our results to the reported activities of ADI-15946 or ADI-23774 (Wec et al., 2017) indicates a high potency for the homologous base-specific antibody rEBOV-515 that we describe. In agreement with this functional assessment, our structural data showed that rEBOV-515 strongly interacts with conserved residues in the GP, with a unique footprint among base-specific broadly reactive human mAbs rEBOV-515, rEBOV-520, and ADI-15946, suggesting a structural basis for its remarkable breadth and potency.

Multifunctionality is another desirable feature in antibody cocktails in addition to neutralizing potency and breadth (Saphire et al., 2018). Reports of human EVD cases revealed high plasma viral RNA titers at the time of patient admission into antibody treatment studies (Brown et al., 2018; Mbala-Kingebeni et al., 2019). Similarly, we observed high serum viral titers in each cohort of challenged NHPs before antibody treatment (**Figure 5**), indicating that at the time of treatment many cells in multiple organs are already infected. This finding highlights the importance to retain Fc-mediated effector function activity in antibody cocktails in order to preserve the ability to eliminate infected cells. In the cocktail of EBOV-442 IgG1 + EBOV-515 LALA-PG, the stronger and broader neutralizing antibody EBOV-515 LALA-PG may act directly to neutralize circulating virus, and this activity is enhanced synergistically in the presence of rEBOV-442 IgG1. Antibody rEBOV-442 IgG1, in addition to direct virus neutralization, may act through Fc-mediated functions by binding to the non-overlapping antigenic site on the GP on infected cells, and this activity could be enhanced reciprocally in the presence of EBOV-515 LALA-PG. Such a combination of activities may facilitate viral clearance and potentially decrease the likelihood of viral mutations that facilitate escape from the cocktail treatment. The relative contribution of these activities to the therapeutic efficacy exhibited by this cocktail needs further investigation.

During acute EVD, circulating infectious virus sometimes seeds immune-privileged tissues, including the brain, eyes, and testes, and persist after clearance from the blood and recovery (Diallo et al., 2016; Subtil et al., 2017; Varkey et al., 2015). Until recently, it was generally assumed that Ebola epidemics start upon zoonotic transmission. On Feb 14, 2021, a new EVD outbreak was declared in Guinea, and viral genome sequencing reports suggested that the outbreak was caused by the Makona variant of EBOV that caused the 2014 EVD epidemics (virological.org, 2021). The index case of the 2021 Guinea cluster likely was infected from a persistent source, such as via sexual transmission from an EVD survivor (virological.org, 2021), raising concerns about possible person-to-person transmission and reignition of outbreaks. The existing therapies and those that are currently in clinical development should be evaluated for their efficacy in clearing infectious virus from immune-privileged sites. Further, NHP studies suggested a genetic drift upon selection pressure with sub-optimal antibody treatment that could be a potential cause for failure during EVD treatment (Kugelman et al., 2015). Potent antibody cocktails like EBOV-442 IgG1 + EBOV-515 LALA-PG may help thwart antigenic drift by targeting non-overlapping vulnerable sites on GP and exhibiting complementary mechanisms of action.

In summary, these studies highlight the power of implementing a rational mAb cocktail development program using structure-function-guided principles (*e.g*., knowledge of binding sites, neutralization breadth, resistance to escape, multifunctionality, synergy etc.). We identified a pan-ebolavirus biologic comprising a two-antibody cocktail that exhibits high efficacy for treatment of all three medically important ebolaviruses with only two doses of mAb. These findings set the stage for clinical evaluation of pan-ebolavirus combination therapy with the two human antibodies rEBOV-442 IgG1 and rEBOV-515 LALA-PG.

## Limitations

The protection of NHPs from advanced EVD using a lower dose of antibody cocktail treatment in a volume that can be delivered by the intramuscular or subcutaneous routes should be assessed in future studies. The developability (manufacturability) of antibodies described here needs to be assessed in future studies. However, the results from our preclinical studies to date suggest favorable physiochemical profiles for these mAbs; both antibodies were stable for >1 month when stored at 4°C and did not aggregate at IgG concentrations >25 mg/mL. Major sequence liabilities were not detected (except a longer than average CDRH3 of rEBOV-442) using online-based Therapeutic Antibody Profiler software (http://opig.stats.ox.ac.uk/webapps/newsabdab/sabpred/tap). Both antibodies occurred naturally in response to ebolavirus infection in human survivors, with natural pairing of the heavy and light chains and isotypes (except LALA-PG modification for rEBOV-515) in their recombinant forms. rEBOV-442 and rEBOV-515 exhibited low or modest autoreactivity to human cells, respectively, as we recently reported (Murin et al., 2021).

## ACKNOWLEDGMENTS

We thank D. Deer for technical assistance with NHP studies. We thank George K. Lewis, Robin Flinko, and Chiara Orlandi for providing EBOV GPkik-293FS eGFP CCR5-SNAP cell line and protocols for RFADCC assay. The Jurkat-EBOV GP cell line was a kind gift from C. Davis and R. Ahmed. This work was supported by U.S. N.I.H. grants U19 AI109711 (to J.E.C. and A.B.), U19 AI142785 (to J.E.C. and T.G.), and U19 AI109762 (ABW), HHS contract HHSN272201400058C (to J.E.C.), and DTRA grant HDTRA1-13-1-0034 (to J.E.C. and A.B.). J.E.C. is a recipient of the 2019 Future Insight Prize from Merck KGaA, which supported this work with a grant. The project was supported by The Vanderbilt Institute for Clinical and Translational Research (VICTR) funded by the National Center for Advancing Translational Sciences (NCATS) Clinical Translational Science Award (CTSA) Program, Award Number 5UL1TR002243-03. The content is solely the responsibility of the authors and does not necessarily represent the official views of the NIH. Work in BSL-4 and ABSL-4 was supported by NIH grant 5UC7AI094660-07 and by the Animal Resource Center of the Galveston National Laboratory.

## AUTHOR CONTRIBUTIONS

P.G., C.D.M., R.W.C., R.C., A.B., T.W.G., A.B.W., and J.E.C. planned the studies. P.G., C.D.M., R.W.C., P.A.I., K.H., V.B., K.N.A., J.B.G., N.K., T.A., R.S.N., R.E.S., N.S., S.J.Z., and R.G.B. conducted experiments. P.G., C.D.M., R.W.C., A.B., T.W.G., A.B.W., and J.E.C. interpreted the studies. P.G., C.D.M., and J.E.C. wrote the first draft of the paper. A.B., T.W.G., A.B.W, and J.E.C. obtained funding. All authors reviewed, edited and approved the paper.

## STAR METHODS

Detailed methods include the following:

KEY RESOURCES TABLE

LEAD CONTACT AND MATERIALS AVAILABILITY

EXPERIMENTAL MODEL AND SUBJECT DETAILS

- Cell lines
- Viruses
- Mouse models
- NHP model

METHOD DETAILS

- Monoclonal antibody production and purification
- GP expression and purification
- ELISA binding assay
- Mammalian cell-surface-displayed GP antibody binding
- Measurement of synergistic GP binding by a combination of antibodies
- Selection of VSV/EBOV GP mutants that escape antibody neutralization
- Neutralization assays
- Measurement of synergistic virus neutralization by a combination of antibodies
- Rapid fluorometric antibody-mediated cytotoxicity assay (RFADCC)
- GP cleavage inhibition
- Mouse challenge
- NHP challenge
- Measurement of virus load in NHP blood or tissues
- NHP serum biochemistry
- Sample preparation for cryogenic electron microscopy
- Cryogenic electron microscopy data collection and processing
- Cryogenic electron microscopy model building and refinement

QUANTIFICATION AND STATISTICAL ANALYSIS

DATA AND CODE AVAILABILITY

### LEAD CONTACT AND MATERIALS AVAILABILITY

Further information and requests for resources and reagents should be directed to and will be fulfilled by the Lead Contact, James E. Crowe, Jr. Materials described in this paper are available for distribution under the Uniform Biological Material Transfer Agreement, a master agreement that was developed by the NIH to simplify transfers of biological research materials.

### EXPERIMENTAL MODEL AND SUBJECT DETAILS

#### Cell lines

Vero-E6 (monkey, female origin) and Vero CCL-81 (monkey, female origin) were obtained from the American Type Culture Collection (ATCC). Vero-E6 cells were cultured in Minimal Essential Medium (MEM) (Thermo Fisher Scientific) supplemented with 10% fetal bovine serum (FBS; HyClone) and 1% penicillin-streptomycin in 5% CO_2_, 37°C. Vero CCL-81 cells were cultured in Dulbecco’s Modified Eagle Medium (DMEM; Thermo Fisher Scientific) supplemented with 10% Ultra-Low IgG FBS (Gibco), 25 mM HEPES, and 100 units/mL of penicillin, and 100 μg/mL of streptomycin (GIBCO) in 5% CO_2_, 37°C. A 293F cell line (human, female origin) stably-transfected to express SNAP-tagged EBOV GP was described previously (Domi et al., 2018). ExpiCHO (hamster, female origin) and FreeStyle 293F (human, female origin) cell lines were purchased from Thermo Fisher Scientific and cultured according to the manufacturer’s protocol. The Jurkat-EBOV GP (Makona variant) cell line stably transduced to display EBOV GP on the surface (Davis et al., 2019) was a kind gift from Carl Davis (Emory University, Atlanta, GA). All cell lines were tested on a monthly basis for *Mycoplasma* and found to be negative in all cases.

#### Viruses

The mouse-adapted EBOV Mayinga variant (EBOV-MA, GenBank: AF49101) (Bray et al., 1998), authentic EBOV Mayinga variant expressing eGFP (Towner et al., 2005), the chimeric infectious EBOV/BDBV-GP (GenBank: <u>KU174137</u>) and EBOV/SUDV-GP (GenBank: <u>KU174142</u>) viruses expressing eGFP (Ilinykh et al., 2016), the infectious vesicular stomatitis viruses rVSV/EBOV GP (Mayinga variant) (Garbutt et al., 2004), rVSV/BDBV GP (Uganda variant) (Mire et al., 2013), or rVSV/SUDV GP (Boniface variant) (Geisbert et al., 2008) expressing an ebolavirus GP that replaces VSV G protein were used for mouse challenge studies or neutralization assays. Viruses were grown and titrated in Vero cell monolayer cultures. Authentic ebolaviruses EBOV (Cross et al., 2016), BDBV (Towner et al., 2008), and SUDV (Thi et al., 2016) were used for NHP challenge studies. The EBOV isolate 199510621 (Kikwit variant) originated from a 65-year-old female patient who died on 5 May 1995. The study challenge material was from the second Vero-E6 passage of EBOV isolate 199510621. The first passage at UTMB consisted of inoculating CDC 807223 (passage 1 of EBOV isolate 199510621) at a MOI of 0.001 onto Vero E6 cells. SUDV isolate 200011676 (variant Gulu) originated from a 35-year-old male patient who died on 16 October 2000. The study challenge material was from the second Vero-E6 cell passage of SUDV isolate 200011676. The first passage at UTMB consisted of inoculating CDC 808892 (CDC passage 1 of SUDV isolate 200011676) at a MOI of 0.001 onto Vero-E6 cells. BDBV isolate 200706291 variant Uganda originated from serum of a patient collected in Uganda on 1 October 2007. The study challenge material was from the second Vero-E6 cell passage of BDBV isolate 200706291 variant Uganda. Briefly, the first passage at UTMB consisted of inoculating CDC 811250 (CDC passage 1 of BDBV isolate 200706291) at a MOI of 0.001 onto Vero-E6 cells. The cell culture fluids were subsequently harvested at day 10 post inoculation with each of indicated viruses and stored at - 80°C as ∼1 mL aliquots. Neither mycoplasma nor endotoxin were detected (˂ 0.5 endotoxin units (EU)/mL).

#### Mouse model

Seven- to eight-week old female BALB/c mice were obtained from the Jackson Laboratory. Mice were housed in microisolator cages and provided food and water *ad libitum*. Challenge studies were conducted under maximum containment in an animal biosafety level 4 (ABSL-4) facility of the Galveston National Laboratory, UTMB. The animal protocols for testing of mAbs in mice were approved by the Institutional Animal Care and Use Committee of the University of Texas Medical Branch (UTMB) in compliance with the Animal Welfare Act and other applicable federal statutes and regulations relating to animals and experiments involving animals.

#### Nonhuman primate (NHP) model

Three- to four-year-old male (n=6) or female (n=6) rhesus macaques used in this study were obtained from PrimGen. Four-year-old male (n=3) or female (n=3) cynomolgus monkeys were obtained from Worldwide Primates. NHP research adhered to principles stated in the eighth edition of the Guide for the Care and Use of Laboratory Animals (National Research Council, 2011). The facility where this research was conducted [University of Texas Medical Branch (UTMB)] is fully accredited by the Association for Assessment and Accreditation of Laboratory Animal Care International and has an approved Office of Laboratory Animal Welfare Assurance (#A3314-01).

### METHOD DETAILS

#### Monoclonal antibody production and purification

Sequences of monoclonal antibodies that had been synthesized as cDNA (Twist Bioscience) and cloned into an IgG1 or IgG1 LALA-PG monocistronic expression vector (designated as pTwist-mCis_G1 or pTwist-mCis_hG1 LALA-PG) were used for monoclonal antibody secretion in mammalian cell culture. This vector contains an enhanced 2A sequence and GSG linker that allows the simultaneous expression of monoclonal antibody heavy and light chain genes from a single construct upon transfection (Chng et al., 2015). CHO cell cultures were transfected using the Gibco ExpiCHO Expression System protocols as described by the vendor. Culture supernatants were purified using 5 mL HiTrap MabSelect SuRe (Cytiva, formerly GE Healthcare Life Sciences) column and an ÄKTA pure chromatography system (Cytiva). Purified monoclonal antibodies were buffer-exchanged into PBS, concentrated using Amicon Ultra-4 50-kDa centrifugal filter units (Millipore Sigma) and stored at 4 °C until use. For NHP treatment studies antibodies were purified from 5 to 15 L of CHO supernatant using HiScale 26/20 column (Cytiva) packed with MabSelect SuRe resin, purified protein was buffer-exchanged into PBS using HiScale 50/40 column packed with Sephadex G-25 (medium) resin (GE Healthcare Life Sciences), concentrated, and stored at -80 °C until use. Purified monoclonal antibodies were tested routinely for endotoxin levels (found to be less than 30 EU per mg IgG for mouse studies and less than 1 EU per mg IgG for NHP studies). Endotoxin testing was performed using the PTS201F cartridge (Charles River), with a sensitivity range from 10 to 0.1 EU per mL, and an Endosafe Nexgen-MCS instrument (Charles River). For structural studies, Fab was produced after co-transfection of ExpiCHO cells with two separate mammalian expression vectors containing antibody light chain and Fab heavy chain sequences as described previously (Gilchuk et al., 2020b). Fab proteins were purified using CaptureSelect column (Thermo Fisher Scientific). Purified antibodies were buffer-exchanged into PBS, concentrated using Amicon Ultra-4 30 kDa MWCO centrifugal filter units (Millipore Sigma) and stored at 4°C until use.

#### GP expression and purification

For ELISA studies, the ectodomains of EBOV GP ΔTM (residues 1-636; strain Makona; GenBank: KM233070), BDBV GP ΔTM (residues 1-643; strain 200706291 Uganda; GenBank: NC_014373), SUDV GP ΔTM (residues 1-637; strain Gulu; GenBank: NC_006432), and MARV GP ΔTM (residues 1-648; strain Angola2005; GenBank: DQ447653) were expressed and purified as described before (Gilchuk et al., 2018a). For structural studies, the ectodomain of EBOV/Makona GP (residues 32-644, GenBank AKG65268.1) lacking residues 310-460 of the mucin-like domain to produce EBOV/Makona GP*Δ*Muc was produced and purified as described before (Murin et al., 2021).

#### ELISA binding assay

To assess mAb binding at different pH, wells of 96-well microtiter plates were coated with purified, recombinant EBOV, BDBV or SUDV GPΔTM ectodomains at 4°C overnight. Plates were blocked with 2% non-fat dry milk and 2% normal goat serum in DPBS containing 0.05% Tween-20 (DPBS-T) for 1 h. Purified mAbs were diluted serially in DPBS-T (pH 7.4), or DPBS-T that was adjusted to pH 5.5 or 4.5 with hydrochloric acid, added to the wells and incubated for 1 h at ambient temperature. The bound antibodies were detected using goat anti-human IgG conjugated with horseradish peroxidase (Southern Biotech) diluted in blocking buffer. Color development was monitored using TMB (3,3′,5,5′-tetramethylbenzidine) substrate (Thermo Fisher Scientific), 1N hydrochloric acid was added to stop the reaction, and the absorbance was measured at 450 nm using a spectrophotometer (Biotek).

#### Mammalian cell-surface-displayed GP antibody binding

Binding of Alexa Fluor 647-labeled antibody to Jurkat-EBOV GP cell line was assessed by flow cytometry using an iQue Screener Plus high throughput flow cytometer (Intellicyt Corp.) as we described previously (Gilchuk et al., 2020b).

#### Measurement of synergistic GP binding by a combination of antibodies

Serially-diluted Alexa Fluor 647-labeled antibody rEBOV-442 IgG1 or rEBOV-515 LALA-PG was titrated into serially-diluted unlabeled partner antibody to generate a pairwise combinatorial matrix of two antibodies in the mixture. For antibody dilutions and washes, we used DPBS (Dulbecco’s phosphate-buffered saline) containing 2% of heat-inactivated FBS and 2 mM EDTA (ethylenediaminetetraacetic acid, sodium salt) (pH 8.0) designated as incubation buffer. For antibody staining, *∼*5 x 10^4^ Jurkat EBOV-GP cells were added per each well of V-bottom 96-well plate (Corning) in 5 µL of the incubation buffer, and antibody mixtures were added to the cells in duplicate for total volume of 50 µL per well, followed by 2 hr incubation at 4°C. Cells were washed with the incubation buffer by centrifugation at 400 x *g* for 5 min at ambient temperature and binding to the GP was assessed using iQue Screener Plus flow cytometer. Data for up to 5,000 events per well were acquired, and data were analyzed with ForeCyt (Intellicyt Corp.) software. Dead cells were excluded from the analysis on the basis of forward and side scatter gate for viable cell population. Binding was calculated as the percent of the maximal median fluorescence intensity signal (MFI) by the highest concentration of respective fluorescently-labeled antibody alone (25 μg/mL). Synergy distribution maps were generated from the dose-response binding matrix using a web application, SynergyFinder 2.0, and data was analyzed using ZIP synergy scoring model (Ianevski et al., 2020).

#### Selection and sequencing of VSV/EBOV GP mutants that escape antibody neutralization

To screen for escape mutations selected in the presence of individual antibodies or antibody cocktails, we used a real-time cell analysis (RTCA) assay and xCELLigence RTCA MP Analyzer (ACEA Biosciences Inc.) with modification of recently described assays (Gilchuk et al., 2020a; Greaney et al., 2021). Fifty (50) μL of cell culture medium (DMEM supplemented with 2% FBS) was added to each well of a 96-well E-plate to obtain a background reading. Eighteen thousand (18,000) Vero cells in 50 μL of cell culture medium were seeded per each well, and plates were placed on the analyzer. Measurements were taken automatically every 15 min and the sensograms were visualized using RTCA software version 2.1.0 (ACEA Biosciences Inc). VSV/EBOV GP virus (20,000 plaque forming units [PFU] per well, ∼1 MOI) was mixed with a saturating neutralizing concentration of individual antibody (10 μg/mL) or two-antibody cocktail (1:1 antibody ratio, 10 μg/mL total antibody concentration) in a total volume of 100 μL and incubated for 1 h at 37°C. At 16-20 h after seeding the cells, the virus-antibody mixtures were added into 8 to 96 replicate wells of 96-well E-plates with cell monolayers. Wells containing only virus in the absence of antibody and wells containing only Vero cells in medium were included on each plate as controls. Plates were measured continuously (every 15 min) for 72 h. The escape mutants were identified by delayed CPE in wells containing antibody. To verify escape from rEBOV-442 antibody selection, isolated viruses were assessed in a subsequent RTCA experiment in the presence of 20 μg/mL of rEBOV-442 or 20 μg/mL of rEBOV-515 or 20 μg/mL of 1:1 cocktail of rEBOV-442+rEBOV-515 (see **Figure S1B**).

To verify escape mutations present in GP protein-expressing VSV antibody-selected escape variants, the escape viruses isolated after RTCA escape screening were propagated in 6-well culture plates with confluent Vero cells in the presence of 20 μg/mL of the rEBOV-442. Viral RNA was isolated using a QiAmp Viral RNA extraction kit (QIAGEN) from aliquots of supernatant containing a suspension of the selected virus population. The GP protein gene cDNA was amplified with a SuperScript IV One-Step RT-PCR kit (Thermo Fisher Scientific) using primers flanking the GP gene. The amplified PCR product (∼2,400 bp) was purified using SPRI magnetic beads (Beckman Coulter) at a 1:1 ratio and sequenced by the Sanger sequence technique using primers giving forward and reverse reads of the glycan cap region of the GP.

#### Neutralization assays

BSL-4 virus neutralization assays were performed using recombinant EBOV-eGFP or chimeric EBOV viruses in which GP was replaced with its counterpart from BDBV or SUDV, as described previously (Ilinykh et al., 2016). Briefly, four-fold dilutions of the respective mAb starting at 200 µg/mL were mixed in triplicate with 400 PFU of the virus in U-bottom 96-well plates and incubated for 1 hr at 37°C. Mixtures were applied on Vero-E6 cell monolayer cultures in 96-well plates and incubated for four days at 37°C. In the absence of mAb neutralizing activity, the infection resulted in uniform eGFP fluorescence from the monolayer of cells that was detected readily by fluorescence microscopy. Fluorescence was measured using Synergy HT microplate reader (BioTek). Half maximal inhibitory concentration (IC_50_) values were determined by nonlinear regression analysis using Prism software.

BSL-2 virus neutralization experiments were performed using the infectious rVSV/EBOV GP, rVSV/BDBV GP, and rVSV/SUDV GP viruses, and we adopted high-throughput RTCA assay that quantify virus-induced cytopathic effect (CPE) (Gilchuk et al., 2020a; Gilchuk et al., 2020b). Viruses were pre-titrated by RTCA to determine dilution of each virus stock to achieve similar CPE kinetics and complete CPE in 32 h after applying virus alone to Vero cells. Fifty (50) μL of cell culture medium (DMEM supplemented with 2% FBS) was added to each well of a 96-well E-plate using a ViaFlo384 liquid handler (Integra Biosciences) to obtain background reading. Eighteen thousand (18,000) Vero cells in 50 μL of cell culture medium were seeded per each well, and the plate was placed on the analyzer. Measurements were taken automatically every 15 min, and the sensograms were visualized using RTCA software version 2.1.0 (ACEA Biosciences Inc). VSV/EBOV GP (∼0.1 MOI, ∼2,000 PFU per well), or VSV/BDBV GP (0.04 MOI, ∼800 PFU per well) or VSV/SUDV GP (0.01 MOI, ∼240 PFU per well) were mixed 1:1 with respective dilution of mAb in triplicate a total volume of 100 μL using DMEM supplemented with 2% FBS as a diluent and incubated for 1 h at 37°C in 5% CO_2_. At 16 to 18 h after seeding the cells, the virus-mAb mixtures were added to the cells in 96-well E-plates. Triplicate wells containing virus only (maximal CPE in the absence of mAb) and wells containing only Vero cells in medium (no-CPE wells) were included as controls. Plates were measured continuously (every 15 min) for 48 h to assess virus neutralization. Normalized cellular index (CI) values at the endpoint (42 h after incubation with the virus) were determined using the RTCA software version 2.1.0 (ACEA Biosciences Inc.). Results were expressed as percent neutralization in the presence of a particular mAb relative to no-CPE control wells minus CI values from control wells with maximum CPE. RTCA IC_50_ values were determined by nonlinear regression analysis using Prism software.

#### Measurement of synergistic virus neutralization by a combination of antibodies

We used RTCA assay to assess neutralizing activity from a pairwise combinatorial matrix of two antibodies in the mixture. Serially-diluted rEBOV-442 (2-fold dilutions) was titrated into serially-diluted rEBOV-515 (two-fold dilutions) and incubated with rVSV/EBOV GP, rVSV/BDBV GP, or rVSV/SUDV GP viruses for 1 h at 37°C. Virus-antibody mixtures were applied to Vero cells grown in 96-well E-plates in duplicates for each virus and using separate plates for each pairwise combinatorial matrix. Triplicate wells containing virus only (maximal CPE in the absence of mAb) and wells containing only Vero cells in medium (no CPE wells) were included as controls. Plates were measured continuously (every 15 minutes) for 48 h to assess virus neutralization. Normalized cellular index (CI) values at the endpoint (42 h after incubation with the virus) were determined using the RTCA software version 2.1.0 (ACEA Biosciences Inc.). Results were expressed as percent neutralization in the presence of a particular mAb relative to control wells with no CPE minus CI values from control wells with maximum CPE. Synergy distribution maps were generated from the dose-response binding matrix using a web application, SynergyFinder 2.0, and data was analyzed using ZIP synergy scoring model (Ianevski et al., 2020).

#### Rapid fluorometric antibody-mediated cytotoxicity assay (RFADCC)

Antibody-dependent cell-mediated cytotoxicity activity of EBOV GP-reactive IgG was quantified with an EBOV-adapted modification of the RFADCC assay (Domi et al., 2018; Orlandi et al., 2016). Briefly, a target cell line was made by transfecting 293F cells with a full-length DNA expressing GP from the EBOV-Kikwit isolate followed by transfecting with two separate DNA constructs expressing eGFP and the chimeric CCR5-SNAP tag protein. The new cell line, designated EBOV GPkik-293FS eGFP CCR5-SNAP, expresses EBOV-Kikwit GP on the plasma membrane, eGFP in the cytoplasm and the SNAP-tag CCR5, which can be specifically labeled with SNAP-Surface AF647 (NEB), on the cell surface (Domi et al., 2018). The unrelated human mAb DENV 2D22 and the Fc effector function disabled mAb rEBOV-515 LALA-PG were used as negative controls for the assay background. The ADCC activity was quantified by incubating three-fold serial dilutions of mAbs with EBOV GPkik-293FS eGFP CCR5-SNAP target cells for 15 min at ambient temperature and then adding human PBMC as effector cells for 2 hrs at 37°C, after which cells were washed once with PBS, fixed with 2% PFA, stained and analyzed using an iQue Screener Plus flow cytometer (Intellicyt Corp.). Data analysis was performed with ForeCyt (Intellicyt Corp.) software. The percentage cytotoxicity of the mAb was determined as the number of target cells losing eGFP signal (by virtue of ADCC) but retaining the surface expression of CCR5-SNAP.

#### GP cleavage inhibition

The assay was performed as we described previously (Gilchuk et al., 2018a). Briefly, Jurkat-EBOV GP cells were pre-incubated with serial dilutions of mAbs in DPBS for 20 min at room temperature, then incubated with thermolysin (Promega) diluted in DPBS to 1 mg/mL for 20 min at 37°C. The reaction was stopped by addition of the incubation buffer containing DPBS, 2% heat-inactivated FBS and 2 mM EDTA (pH 8.0). Washed cells were incubated with 5 μg/mL of Alexa Fluor 647-labeled EBOV GP RBD-reactive mAb MR78 (Bornholdt et al., 2016) at 4°C for 60 min. Stained cells were washed, fixed, and analyzed by flow cytometry using Intellicyt iQue. Cells were gated for the viable population, and median fluorescence intensity from Alexa Fluor 647 was determined. Background staining was determined from binding of the labeled mAb MR78 to Jurkat-EBOV GP (uncleaved) cells. Results are expressed as the percent of RBS exposure inhibition in the presence of tested mAb relative to controls for minimal binding of labeled MR78 mAb-only to intact (uncleaved) Jurkat EBOV-GP, and maximal binding of labeled MR78 mAb-only to cleaved Jurkat-EBOV GP.

#### Mouse challenge

Groups of mice (n = 5 per group) were inoculated with 1,000 PFU of the EBOV-MA by the intraperitoneal (i.p.) route. Mice were treated i.p. with indicated doses of individual mAbs on 1 day after virus inoculation (dpi). Human mAb DENV 2D22 served as a control. Mice were monitored twice daily from 0 to 14 dpi for illness, survival, and weight loss, followed by once daily monitoring from 15 dpi to the end of the study at 28 dpi. The extent of disease was scored using the following parameters: score 1 – healthy; score 2 – ruffled fur and hunched posture; score 3 – a score of 2 plus one additional clinical sign such as orbital tightening and/or >15% weight loss; score 4 – a score of 3 plus one additional clinical sign such as reluctance to move when stimulated, or any neurologic signs (seizures, tremors, head tilt, paralysis, etc.), or >20% weight loss. Animals reaching a score of 4 were euthanized as per the IACUC-approved protocol. All mice were euthanized on day 28 after EBOV challenge.

#### NHP challenge

Twelve healthy adult rhesus macaques (*Macaca mulatta*) of Chinese origin and six healthy adult cynomolgus monkeys (*Macaca fascicularis*) were studied. Animals were assigned to three groups of five animals per treatment group and a control untreated animal. Animals were randomized by random number assignment (with Microsoft Excel) into a treatment group and a control animal. After intramuscular challenge with a lethal target dose of 1,000 plaque-forming units (PFU) of EBOV/Kikwit, BDBV/Uganda, or SUDV/Gulu, each of the NHPs of the treatment group received intravenously two 30 mg/kg doses of the cocktail (a 2:1 mixture of rEBOV-515 LALA-PG and rEBOV-442 IgG1) spaced 3 days apart (days 3 and 6 after EBOV/Kikwit, days 6 and 9 after BDBV/Uganda, or days 4 and 7 after SUDV SUDV/Gulu inoculation). The back-titer of the EBOV, BDBV, and SUDV inoculum identified 963, 1113, and 988 PFU as the actual inoculation dose for the respective virus. Historical untreated controls for EBOV challenge cohort included fourteen untreated animals from separate studies including 11 animals, as we reported previously (Gilchuk et al., 2020b), which were challenged with the same target dose of EBOV/Kikwit and by the same route. Historical untreated controls for SUDV/Gulu challenge cohort included five untreated animals from previous study (Thi et al., 2016) and four untreated animals from two other studies (Geisbert and Cross, unpublished) that were challenged with the same target dose of BDBV and by the same route. Historical untreated controls for BDBV challenge cohort included three untreated animals from a previous study (Bornholdt et al., 2019) and seventeen untreated animals from the other studies (Geisbert and Cross, unpublished) that were challenged with the same target dose of BDBV/Uganda and by the same route. All animals were given physical exams, and blood was collected at the time of inoculation and at indicated times after virus inoculation. In addition, all animals were monitored daily and scored for disease progression with an internal filovirus scoring protocol approved by the UTMB Institutional Animal Care and Use Committee. The scoring measured from baseline and included posture or activity level, attitude or behavior, food and water intake, respiration, and disease manifestations such as visible rash, hemorrhage, ecchymosis, or flushed skin. A score of ≥ 9 indicated that an animal met criteria for euthanasia. These studies were not blinded, and all animals were included in analysis.

#### Measurement of virus load in NHP blood and tissues

Titration of virus in plasma samples and 10% tissue homogenates (w/v) was performed by plaque assay in Vero-E6 cell culture monolayers. Briefly, serial 10-fold dilutions of the samples were applied to Vero-E6 cell monolayers in duplicate wells (200 µL); the limit of detection was 25 PFU/mL for plasma and 250 PFU/gram for tissue. For qRT-PCR analysis, RNA was isolated from whole blood or tissue using the Viral RNA Mini-kit (Qiagen) using 100 µL of blood or 100 mg of tissue into 600 µL of buffer AVL. Primers (probes) targeting the VP30 gene of EBOV probe sequence of 6-carboxyflourescein (6FAM)-5*′* CCG TCA ATC AAG GAG CGC CTC 3*′*-6 carboxytetramethylrhodamine (TAMRA) (Thermo Fisher Scientific), the VP35 intergenic region of BDBV probe sequence of 6FAM-5*′*CGCAACCTCCACAGTCGCCT 3*′*-TAMRA, and the L gene of SUDV probe sequence of 6FAM-5’ CAT CCA ATC AAA GAC ATT GCG A 3*′*-TAMRA were used for qRT-PCR. EBOV RNA was detected using the CFX96 detection system (BioRad Laboratories) in One-step probe qRT-PCR kits (Qiagen) with the following cycle conditions: 50 °C for 10 min, 95 °C for 10 s, and 40 cycles of 95 °C for 10 s and 57 °C for 30 s for EBOV and BDBV and 50 °C for 10 min, 95 °C for 10 s, and 40 cycles of 95 °C for 10 s and 59 °C for 30 s for SUDV. Threshold cycle (CT) values representing EBOV, BDBV and SUDV genomes were analyzed with CFX Manager Software, and data are depicted as genome equivalents (GEq); the limit of detection was 3.7 log_10_GEq/mL for blood and 3.7 log_10_GEq/g for the tissues.

#### NHP serum biochemistry

Serum samples collected from NHPs were tested for concentrations of albumin, amylase, alanine aminotransferase, aspartate aminotransferase, alkaline phosphatase, gamma-glutamyltransferase, glucose, blood urea nitrogen, creatinine, total protein, and C-reactive protein by using a Piccolo point-of-care analyzer and Biochemistry Panel Plus analyzer discs (Abaxis).

#### Sample preparation for cryogenic electron microscopy

EBOV/Makona GPΔmuc was incubated overnight with a 5-fold molar excess of rEBOV-515 Fab, rEBOV-442 Fab, and rADI-16061 Fab at 4°C. The complexes were then purified by SEC using an S200I column equilibrated in 1X TBS and concentrated to 5 mg/mL using a 100-kDa concentrator (Amicon Ultra, Millipore). Immediately prior to freezing, 0.06 mM of n-Dodecyl *β*-D-maltoside (Anatrace) was added to 3 µL of the complex. Vitrification was performed with a Vitrobot (Thermo Fisher Scientific) equilibrated to 4°C and 100% humidity. Cryo-EM grids were plasma cleaned for 5 s using a mixture of Ar/O2 (Gatan Solarus 950 Plasma system) followed by blotting on both sides of the grid with filter paper (Whattman No. 1). See Table S7 for additional details. Note that ADI-16061 Fab was added to assist in angular sampling and orientations of the complexes in ice as we described previously (Gilchuk et al., 2020b).

#### Cryogenic electron microscopy data collection and processing

Cryo-EM data were collected according to Table S1. Micrographs were aligned and dose-weighted using MotionCor2 (Zheng et al., 2017). CTF estimation was completed using GCTF (Zheng et al., 2017). Particle picking and initial 2D classification were initially performed using CryoSPARC 2.0 (Punjani et al., 2017) to clean up particle stacks and exclude any complexes that were degrading. Particle picks were then imported into Relion 3.1 (Zivanov et al., 2018) for 3D classification and refinement using C3 symmetry and a tight mask around the GP/rEBOV-515 Fab/rEBOV-442 Fab complex. CTF refinement was then performed by either Relion or Cryosparc to increase map quality and resolution. There was no electron density for ADI-16061 Fab.

#### Cryogenic electron microscopy model building and refinement

Homology models of Fab were first generated using SWISS-MODEL (Biasini et al., 2014). A model of EBOV GP (PDB: 5JQ3) was then added to generate a starting model used for refinement. The starting model was fit into the cryo-EM map using UCSF Chimera (Pettersen et al., 2004) and refined initially using Phenix real-space refinement (Liebschner et al., 2019). The refined model was then used as a starting model for relaxed refinement in Rosetta (DiMaio et al., 2015). The top five models then were evaluated for fit in EM density and adjusted manually using Coot (Emsley et al., 2010) to maximize fit. Finally, Man9 glycans were fit into glycan densities, trimmed and then a final refinement was performed in Rosetta. The final structures were evaluated using EMRinger (Barad et al., 2015) and Molprobity from Phenix. Glycans were validated using Privateer (Agirre et al., 2015) and PDBcare (Lutteke and von der Lieth, 2004). All map and model images were generated in UCSF Chimera (Pettersen et al., 2004). Antibody contacts were analyzed using LigPlot (Laskowski and Swindells, 2011), Arpeggio (Jubb et al., 2017) and UCSF Chimera (Pettersen et al., 2004).

### QUANTIFICATION AND STATISTICAL ANALYSIS

The descriptive statistics mean ± SEM or mean ± SD were determined for continuous variables as noted. Survival curves were estimated using the Kaplan-Meier method and overall difference in survival between the groups in mouse studies was estimated using two-sided log rank test (Mantel-Cox) with subjects right censored, if they survived until the end of the study. In NHP studies, survival curves were estimated using the Kaplan-Meier method, and the proportion surviving at day 28 after virus inoculation was compared using a 2-sided exact unconditional test of homogeneity. Curves for antibody binding were fitted after log transformation of antibody concentrations using a four-parameter log-logistic (4PL) analysis. In neutralization assays and GP cleavage inhibition assays, IC_50_ values were calculated after log transformation of antibody concentrations using a four-parameter log-logistic (4PL) analysis. Synergy distribution maps were generated from the dose-response binding matrix using an open-source software SynergyFinder 2.0: visual analytics of multi-drug combination synergies (https://synergyfinder.fimm.fi), and data was analyzed using ZIP synergy scoring model (Ianevski et al., 2020). Technical and biological replicates are indicated in the figure legends. Statistical analyses were performed using Prism v8.4.3 (GraphPad).

### DATA AND CODE AVAILABILITY

The EBOV GP ΔMuc ΔTM (Makona)*−*rEBOV-515*−*rEBOV-442 Fab cryo-EM structure has been deposited in the PDB with accession code 7M8L. The accession number for cryo-EM reconstructions reported in this paper have been deposited to the Electron Microscopy Data Bank under accession EMDB code EMD-23719 (see Key Resources Table for details). All data needed to evaluate the conclusions in the paper are present in the paper or the Supplemental Information; source data for each of the display items is provided in Key Resources Table.

