## Supplemental Figures and Tables for "Protective pan-ebolavirus combination therapy by two multifunctional human antibodies"

Figure S1

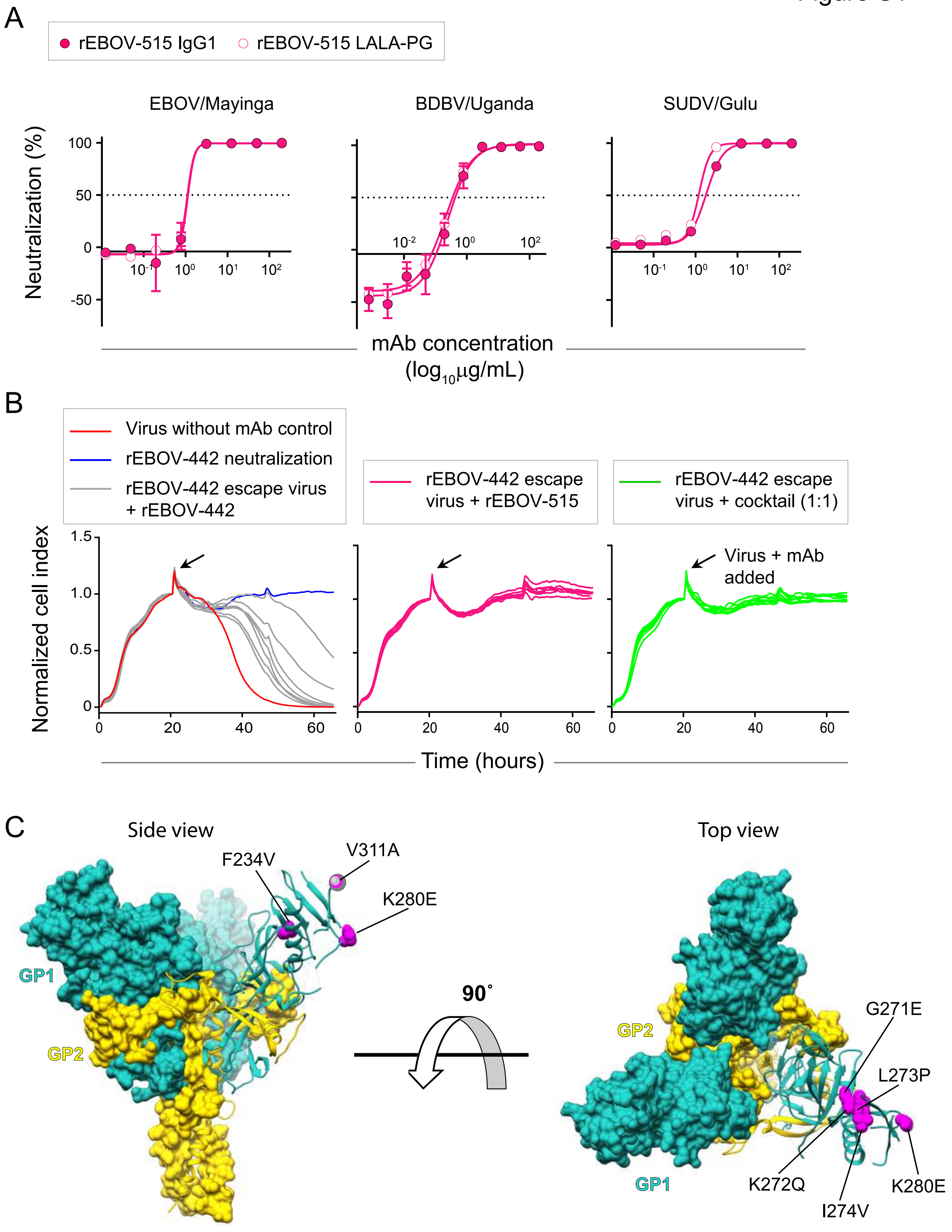
A
rEBOV-515 IgG1
rEBOV-515 LALA-PG
EBOV/Mayinga
BDBV/Uganda
SUDV/Gulu
Neutralization (%)
100
50
0
-50
$10^{-2}$
$10^{-1}$
$10^{0}$
$10^{1}$
$10^{2}$
mAb concentration
($\log_{10}\mu g/mL$)
B
Virus without mAb control
rEBOV-442 neutralization
rEBOV-442 escape virus + rEBOV-442
rEBOV-442 escape virus + rEBOV-515
rEBOV-442 escape virus + cocktail (1:1)
Virus + mAb added
Normalized cell index
1.5
1.0
0.5
0
20
40
60
Time (hours)
C
Side view
Top view
F234V
V311A
K280E
GP1
GP2
90˚
G271E
L273P
K272Q
I274V

**Figure S1**. **Neutralizing activity of antibody rEBOV-515 LALA-PG and neutralization of antibody rEBOV-442-selected escape viruses by rEBOV-515 or the cocktail.**

(A) EBOV, BDBV, or SUDV neutralization by wild-type IgG1 or IgG1 LALA-PG variants of rEBOV-515. Neutralization was assessed with chimeric Ebola viruses encoding EBOV, DBDV, or SUDV GP as in **Figure 1A**. Mean ± SD of technical triplicates from one experiment are shown.

(B) Real-time cell analysis (RTCA) sensograms for antibody rEBOV-442-selected escape viruses for neutralization by rEBOV-442 alone, rEBOV-515 alone, or a 1:1 antibody cocktail. The escape selection was performed using infectious chimeric VSV expressing EBOV (Mayinga variant) GP. Seven viruses that escaped neutralization by rEBOV-442 and had sequence-verified mutations in the gene encoding the glycan cap region of the GP were incubated in the presence of a saturating concentration (20 μg/mL) of antibody rEBOV-442 (left panel, grey), rEBOV-515 (middle, rose), or a 1:1 mixture of both antibodies. Cytopathic effect (CPE) was monitored kinetically in Vero cells after applying the virus-antibody mixtures to the cells. Sensograms for CPE by the wild-type VSV/EBOV GP without antibody (left panel, red), and full neutralization of the wild-type VSV/EBOV GP by rEBOV-442 (left panel, blue) are shown as the controls. Seven viruses with individual mutations selected by rEBOV-442 and indicated in **Figure 1D**, escape neutralization by rEBOV-442 (left panel, grey) but were neutralized by rEBOV-515 (middle panel) or a 1:1 mixture of rEBOV-442 and rEBOV-515 (right panel).

(C) Mutations in indicated residues (magenta) identified in the glycan cap region of the GP (PDB: 5JQ3) for rEBOV-442-selected viruses that escaped neutralization by rEBOV-442 as in (B). Position of V311 residue is shown with grey sphere.

Related to **Figure 1**.

Figure S2

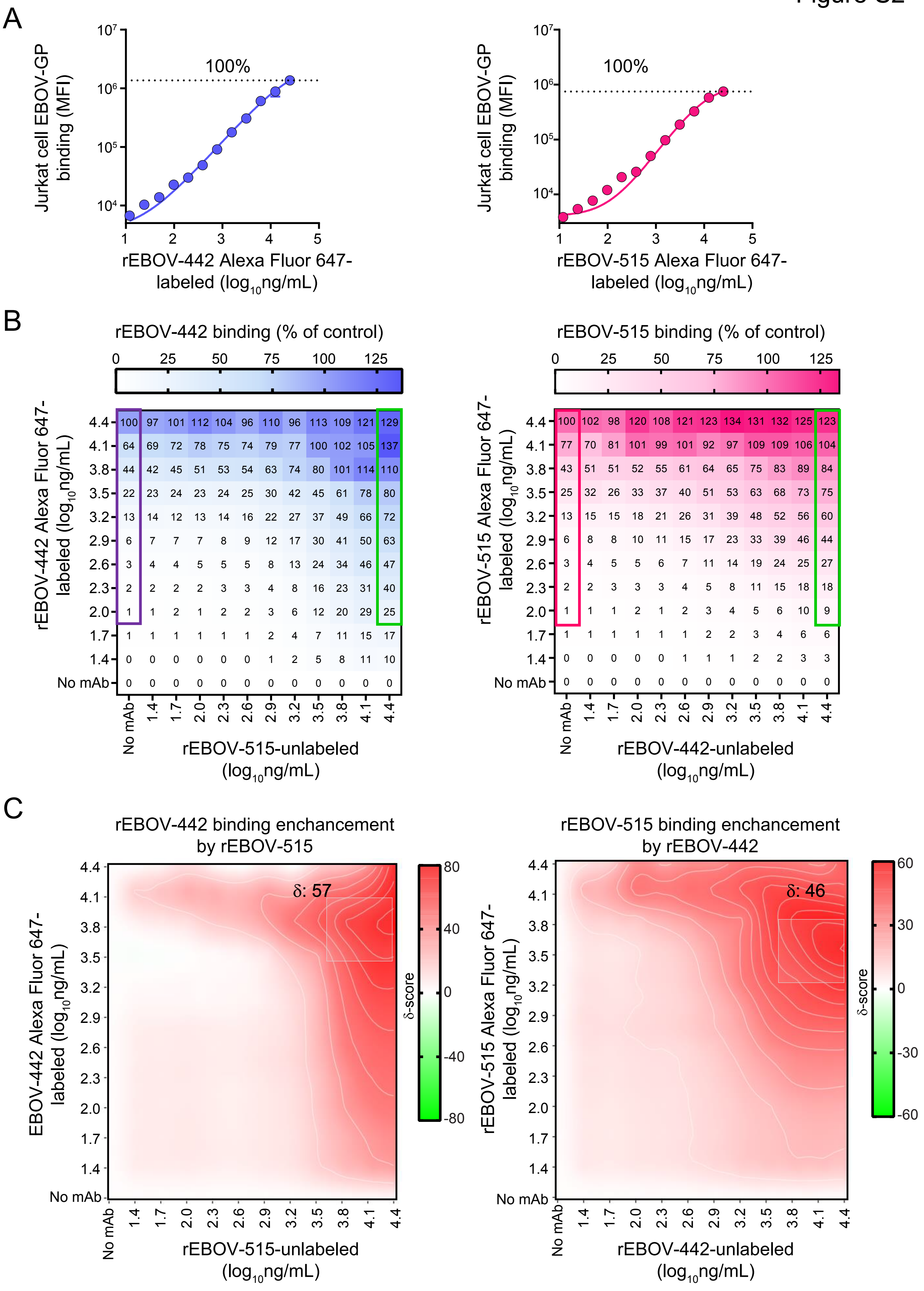
A
Jurkat cell EBOV-GP binding (MFI)
100%
rEBOV-442 Alexa Fluor 647-labeled (log10ng/mL)
rEBOV-515 Alexa Fluor 647-labeled (log10ng/mL)
B
rEBOV-442 binding (% of control)
0 25 50 75 100 125
rEBOV-442 Alexa Fluor 647-labeled (log10ng/mL)
4.4: 100 97 101 112 104 96 110 96 113 109 121 129
4.1: 64 69 72 78 75 74 79 77 100 102 105 137
3.8: 44 42 45 51 53 54 63 74 80 101 114 110
3.5: 22 23 24 23 24 25 30 42 45 61 78 80
3.2: 13 14 12 13 14 16 22 27 37 49 66 72
2.9: 6 7 7 7 8 9 12 17 30 41 50 63
2.6: 3 4 4 5 5 5 8 13 24 34 46 47
2.3: 2 2 2 2 3 3 4 8 16 23 31 40
2.0: 1 1 2 1 2 2 3 6 12 20 29 25
1.7: 1 1 1 1 1 1 2 4 7 11 15 17
1.4: 0 0 0 0 0 0 1 2 5 8 11 10
No mAb: 0 0 0 0 0 0 0 0 0 0 0 0
No mAb 1.4 1.7 2.0 2.3 2.6 2.9 3.2 3.5 3.8 4.1 4.4
rEBOV-515-unlabeled (log10ng/mL)
rEBOV-515 binding (% of control)
0 25 50 75 100 125
rEBOV-515 Alexa Fluor 647-labeled (log10ng/mL)
4.4: 100 102 98 120 108 121 123 134 131 132 125 123
4.1: 77 70 81 101 99 101 92 97 109 109 106 104
3.8: 43 51 51 52 55 61 64 65 75 83 89 84
3.5: 25 32 26 33 37 40 51 53 63 68 73 75
3.2: 13 15 15 18 21 26 31 39 48 52 56 60
2.9: 6 8 8 10 11 15 17 23 33 39 46 44
2.6: 3 4 5 5 6 7 11 14 19 24 25 27
2.3: 2 2 3 3 3 4 5 8 11 15 18 18
2.0: 1 1 1 2 1 2 3 4 5 6 10 9
1.7: 1 1 1 1 1 1 2 2 3 4 6 6
1.4: 0 0 0 0 0 1 1 1 2 2 3 3
No mAb: 0 0 0 0 0 0 0 0 0 0 0 0
No mAb 1.4 1.7 2.0 2.3 2.6 2.9 3.2 3.5 3.8 4.1 4.4
rEBOV-442-unlabeled (log10ng/mL)
C
rEBOV-442 binding enchancement by rEBOV-515
δ: 57
EBOV-442 Alexa Fluor 647-labeled (log10ng/mL)
rEBOV-515-unlabeled (log10ng/mL)
δ-score
80 40 0 -40 -80
rEBOV-515 binding enchancement by rEBOV-442
δ: 46
rEBOV-515 Alexa Fluor 647-labeled (log10ng/mL)
rEBOV-442-unlabeled (log10ng/mL)
δ-score
60 30 0 -30 -60

**Figure S2. Reciprocal synergistic binding activity in the cocktail of rEBOV-442 and rEBOV-515**. Serially-diluted Alexa Fluor 647-labeled antibody rEBOV-442 IgG1 or rEBOV-515 LALA-PG was titrated into serially-diluted unlabeled partner antibody to generate a pairwise combinatorial matrix of two antibodies in the mixture. Binding to Jurkat EBOV-GP cells was assessed by flow cytometric analysis.

(A) Dose-response binding of fluorescently-labeled rEBOV-442 alone (left) or rEBOV-515 alone (right) to Jurkat-EBOV GP. Mean ± SD of technical duplicates is shown.

(B) Binding dose-response matrix by rEBOV-442 or rEBOV-515 bound to the EBOV-GP in the presence of a partner antibody. Axes denote the concentration of each antibody, with the percent binding shown in each square. Binding was calculated as the percent of the maximal median fluorescence intensity signal (MFI) by the highest concentration of respective fluorescently-labeled antibody alone (25 μg/mL) as in (A). Examples of an enhanced binding activity by a combination of fluorescently-labeled antibody plus partner antibody at the highest tested concentration (green box) in a comparison to the binding by fluorescently-labeled rEBOV-442 alone (violet box) or rEBOV-515 alone (raspberry box) are shown.

(C) Synergy distribution map generated from the dose-response binding matrix in (B). Red color indicates areas in which synergistic neutralization was observed; shaded grey box indicates the area of maximum synergy between the two monoclonal antibodies, and $\delta$-score for this area is shown. $\delta$-score is a synergy score: a value of -10 indicates antagonism; a value of -10 to 10 indicates an additive effect; a value of >10 indicates synergy. Data in (A-B) are from a representative experiment performed in technical duplicate and repeated twice.

Related to **Figure 3**.

### Figure S3

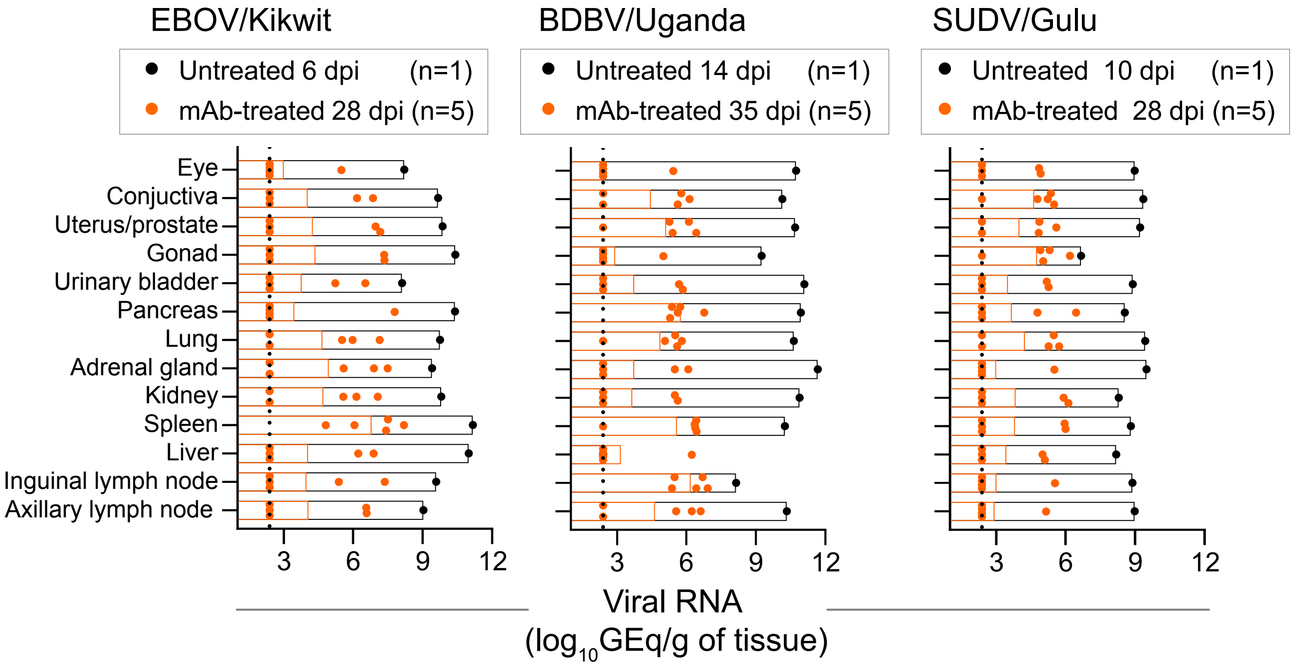
EBOV/Kikwit
Untreated 6 dpi (n=1)
mAb-treated 28 dpi (n=5)
BDBV/Uganda
Untreated 14 dpi (n=1)
mAb-treated 35 dpi (n=5)
SUDV/Gulu
Untreated 10 dpi (n=1)
mAb-treated 28 dpi (n=5)
Eye
Conjuctiva
Uterus/prostate
Gonad
Urinary bladder
Pancreas
Lung
Adrenal gland
Kidney
Spleen
Liver
Inguinal lymph node
Axillary lymph node
3
6
9
12
Viral RNA
(log10GEq/g of tissue)

**Figure S3. The cocktail treatment provides pan-ebolavirus protection of nonhuman primates against viremia.** Viral RNA load in various peripheral tissues of treated NHPs determined by qRT-PCR (28 dpi for EBOV and SUDV cohort, or 35 dpi for BDBV cohort). Tissues from succumbed untreated NHP from each cohort used as a control. The black dotted line indicates the limit of detection, which was 2.4 $log_{10}$GEq/mL. The data are shown as an aligned dot plot with bar, where orange dots indicate measurement from individual treated animals, black dots and bars indicate measurement from untreated control animals, and orange bar indicates the mean value of the measurements for treated cohorts. For samples in which viral RNA was not detected, the measurement values were set to the limit of detection 2.4 $log_{10}$GEq/mL. Each measurement represents the mean of technical duplicates.

Related to **Figure 5**.

Figure S4

A

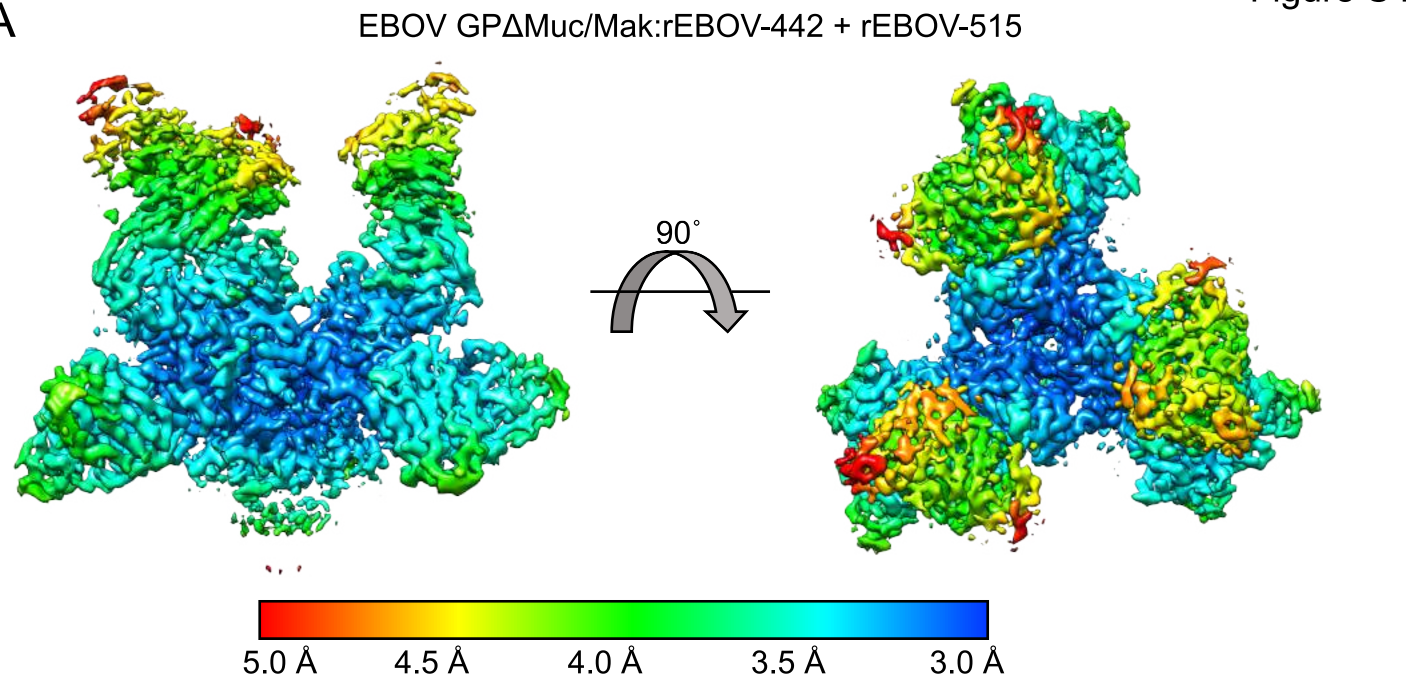

EBOV GPΔMuc/Mak:rEBOV-442 + rEBOV-515
90˚
5.0 Å
4.5 Å
4.0 Å
3.5 Å
3.0 Å


B

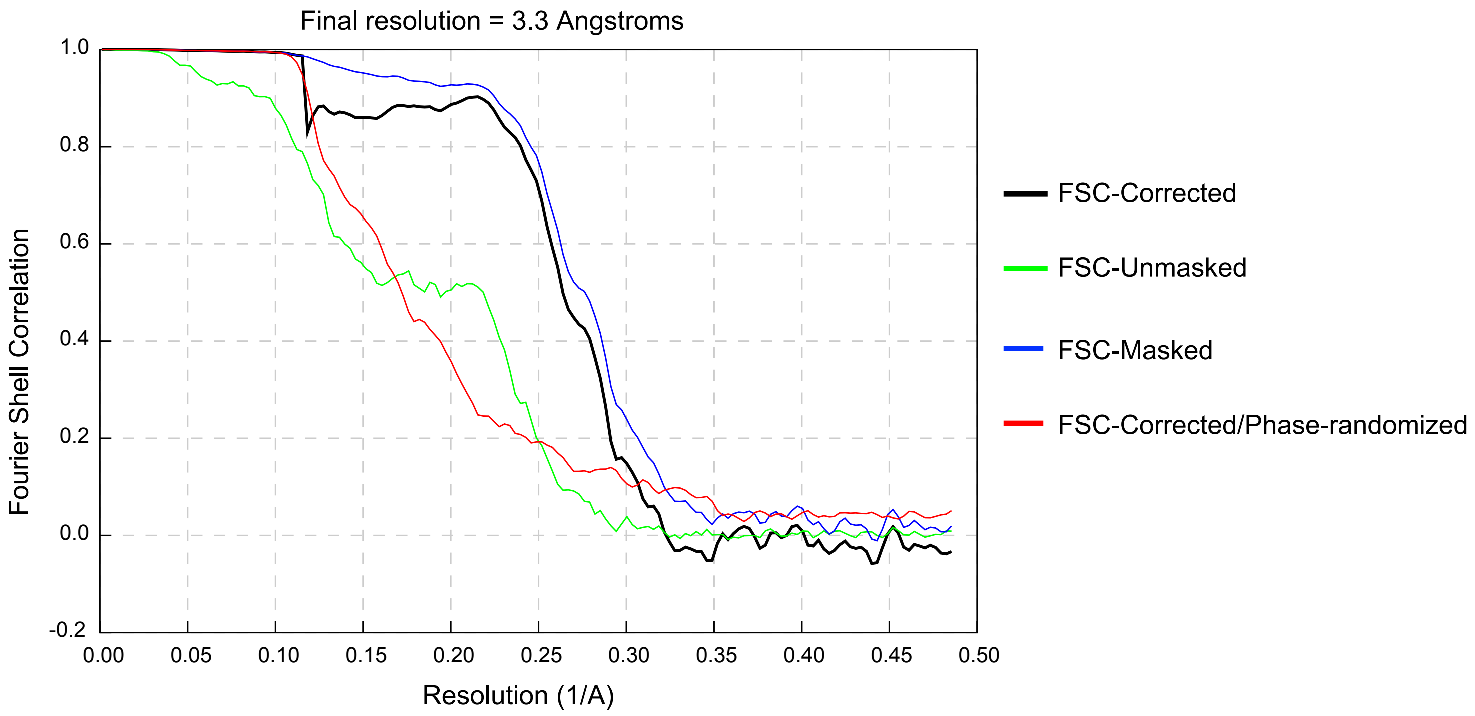

Final resolution = 3.3 Angstroms
Fourier Shell Correlation
Resolution (1/A)
1.0
0.8
0.6
0.4
0.2
0.0
-0.2
0.00
0.05
0.10
0.15
0.20
0.25
0.30
0.35
0.40
0.45
0.50
FSC-Corrected
FSC-Unmasked
FSC-Masked
FSC-Corrected/Phase-randomized

**Figure S4. Cryo-EM local resolution estimation and FSC curves.** (A) The local resolution of the EBOV GPΔMuc/Mak:rEBOV-442/rEBOV-515 cryo-EM map, with local resolutions indicated according to the colors in the legend below. (B) The Fourier shell correlation (FSC) curve for the EBOV GPΔMuc/Mak:rEBOV-442/rEBOV-515 cryo-EM map, generated by Relion 3.1.

Related to **Figure 6**.

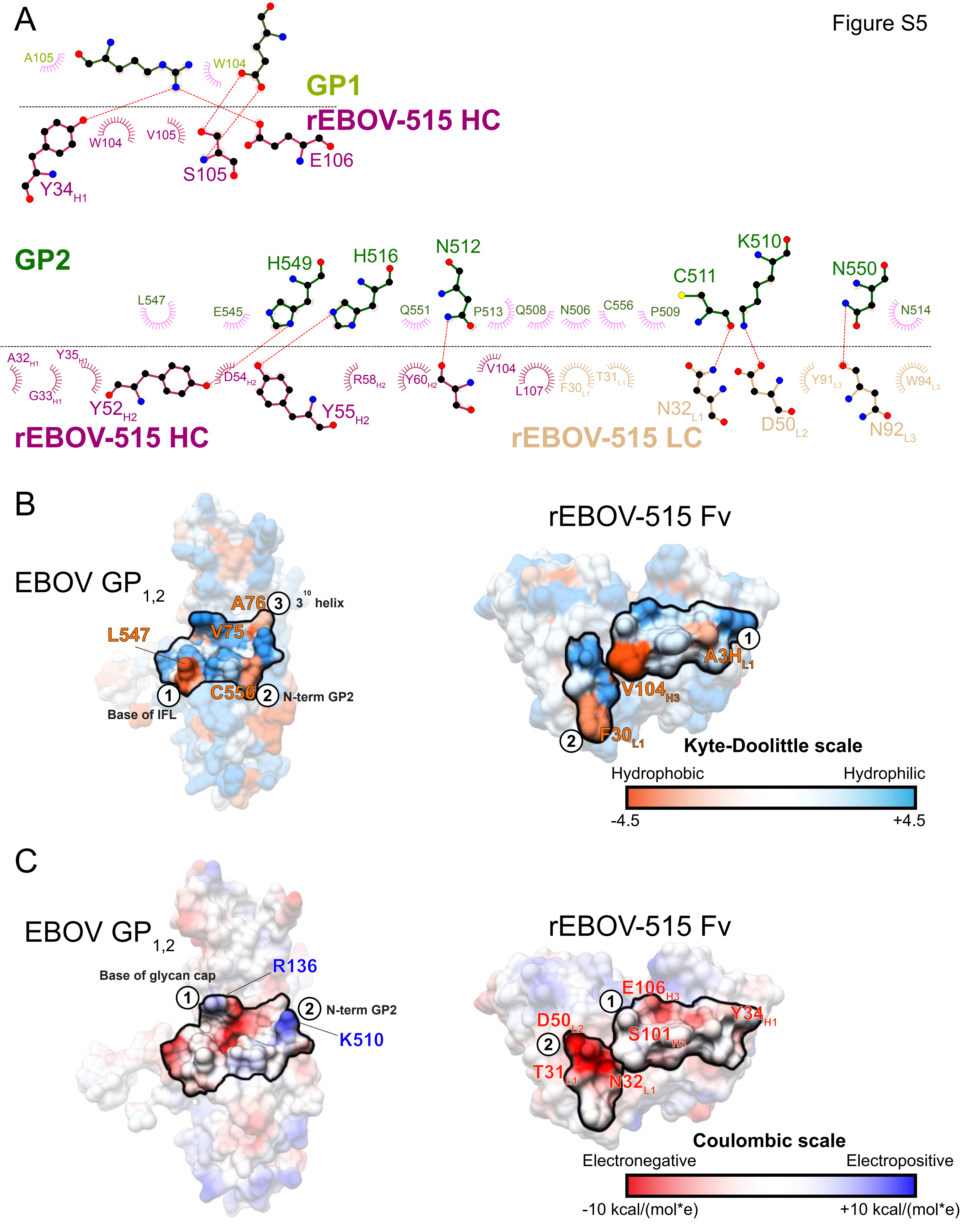

A
Figure S5
A105
W104
GP1
rEBOV-515 HC
W104
V105
S105
E106
Y34H1
GP2
L547
E545
H549
H516
N512
Q551
P513
Q508
N506
C556
P509
C511
K510
N550
N514
A32H1
Y35H1
G33H1
Y52H2
D54H2
Y55H2
R58H2
Y60H2
V104
L107
F30L1
T31L1
N32L1
D50L2
Y91L3
W94L3
N92L3
rEBOV-515 HC
rEBOV-515 LC
B
EBOV GP1,2
A76
3
3 10 helix
L547
V75
1
Base of IFL
C556
2
N-term GP2
rEBOV-515 Fv
1
A3HL1
V104H3
2
F30L1
Kyte-Doolittle scale
Hydrophobic
Hydrophilic
-4.5
+4.5
C
EBOV GP1,2
Base of glycan cap
R136
1
2
N-term GP2
K510
rEBOV-515 Fv
E106H3
1
Y34H1
D50L2
2
S101H3
T31L1
N32L1
Coulombic scale
Electronegative
Electropositive
-10 kcal/(mol*e)
+10 kcal/(mol*e)

**Figure S5. Epitope and paratope binding properties of rEBOV-515.**

(A) LigPlot schematic showing interaction of rEBOV-515 with GP1 (above) or GP2 (below). Residues that make hydrogen bonds are rendered, and those that contribute van der Waals interactions are drawn as semi-circles. rEBOV-515 CDR loop residues are colored according to the labels, with GP residues in magenta/pink and backbone residues colored in dark cyan. Chain identifiers are in parenthesis.

(B) Kyte-Doolittle surface rendering of $GP_{1,2}$ (left) and a face-on view of the rEBOV-515 Fv paratope (right) with the interacting regions highlighted. Key hydrophobic residues are indicated with important regions of interaction labeled 1-3 on the GP epitope and the corresponding interacting residues labeled 1-3 on the rEBOV-515 paratope.

(C) Similar epitope/paratope interaction rendering as in (C) but with Coulombic surface rendering, indicating key electronegative regions on GP that interact with electropositive regions on the rEBOV-515 paratope.

Related to **Figure 6**.

Figure S6

| Subunit | Residue number | Total Contacts | Residue | rEBOV-515 | rEBOV-520 | ADI-15946 |
|---|---|---|---|---|---|---|
| **GP1** | 74 | 8 | G | > | 6 | 2 |
| | 75 | 4 | V | > | 4 | > |
| | 77 | 2 | T | > | 2 | > |
| | 78 | 3 | D | > | 3 | > |
| | 104 | 1 | W | 1 | > | > |
| | 106 | 28 | E | 12 | 5 | 11 |
| | 107 | 7 | N | > | 7 | > |
| | 136 | 29 | R | 13 | 5 | 11 |
| | 137 | 2 | Y | > | 2 | > |
| | 250 | 3 | P | > | 3 | > |
| | 251 | 1 | Q | > | 1 | > |
| | 255 | 5 | Q | > | 5 | > |
| **GP2** | 506 | 8 | N | 5 | > | 3 |
| | 508 | 2 | Q | 1 | > | 1 |
| | 509 | 2 | P | 1 | > | 1 |
| | 510 | 24 | K | 4 | 5 | 15 |
| | 511 | 14 | C | 4 | 3 | 7 |
| | 512 | 29 | P | 3 | 14 | 12 |
| | 513 | 14 | N | 6 | 3 | 5 |
| | 514 | 22 | P | 6 | 5 | 11 |
| | 516 | 9 | H | 5 | 2 | 2 |
| | 545 | 3 | E | 3 | > | > |
| | 547 | 24 | L | 13 | 11 | > |
| | 549 | 20 | H | 11 | 3 | 6 |
| | 550 | 2 | N | 1 | > | 1 |
| | 556 | 2 | C | 2 | > | > |

**Figure S6. Contact residues for broadly reactive antibodies rEBOV-515, rEBOV-520, and ADI-15946**

Base binding antibody contacts on EBOV GP determined by UCSF Chimera with a default cutoff value of -0.4 Å and an allowance of 0 Å (Pettersen et al., 2004). rEBOV-520 from PDB 6PCI (Gilchuk et al., 2020b), ADI-15946 from PDB 6MAM (West et al., 2019). Gradient is green to red, with green being fewer contacts and red being the most contacts.

Related to **Figure 6**.

Figure S7

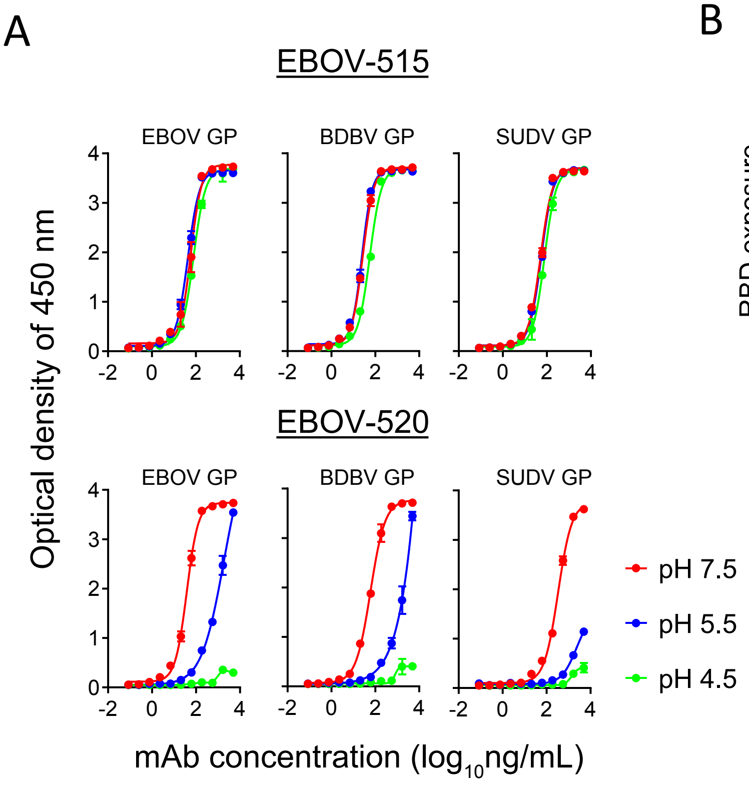


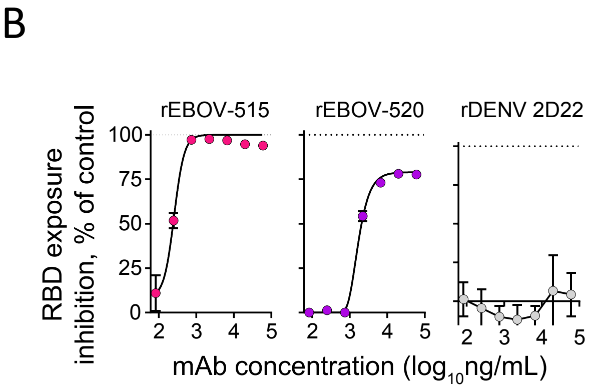


| mAb | $IC_{50}$ (μg/mL) | $IC_{90}$ (μg/mL) |
|---|---|---|
| rEBOV-515 | 0.3 | 0.5 |
| rEBOV-520 | 2.1 | > |
| rDENV 2D22 | > | > |

**Figure S7. rEBOV-515 retains GP-binding capacity at low pH and efficiently inhibits GP cleavage.**

(A) ELISA binding of rEBOV-515 or rEBOV-520 to the recombinant EBOV, BDBV, or SUDV GP ΔTM at neutral or acidic pH. The mean ± SD of technical triplicates and one of two independent experiments are shown.

(B) Cleavage inhibition by the GP-base specific antibodies. Jurkat EBOV-GP cells were pre-incubated with various concentrations of indicated antibodies, treated with thermolysin, then incubated with fluorescently labeled antibody MR78 that recognizes the receptor binding site (RBS) of the GP and assessed for exposure of the RBS on cleaved GP by flow cytometry. Antibody BDBV223 that target a distal antigenic site in the canonical heptad repeat 2 (HR2) region near the membrane proximal external region (MPER) of GP, and dengue virus-specific antibody rDENV 2D22 were used as controls. Results are expressed as the percent of RBS exposure inhibition in the presence of tested mAb relative to controls for minimal binding of labeled MR78 mAb-only to intact (uncleaved) Jurkat EBOV-GP, and maximal binding of labeled MR78 mAb-only to cleaved Jurkat-EBOV GP. Mean ± SD of technical triplicates and one of two independent experiments are shown.

Related to **Figure 6**.

**Table S1. Hematology measurements from rEBOV-515+rEBOV-442 cocktail-treated or -untreated individual NHP that were challenged with EBOV. Related to Figure 4**

| Animal ID | Day after challenge | WBC per mm$^3$ ( x 10$^3$) | RBC per mm$^3$ (x 10$^3$) | HgB (g/dL) | Hct (%) | Platelets per mm$^3$ (x 10$^3$ | Lymph per mm$^3$ (x 10$^3$) | Lymph. (%) | Mono. per mm$^3$ (x 10$^3$) | Neutr. (%) | Eosin. (%) | Baso. (%) | MCH (pg/cell) | MPV (fL/cell) |
|---|---|---|---|---|---|---|---|---|---|---|---|---|---|---|
| M1 | 0 | 8 | 5 | 11 | 37 | 326 | 3 | 30 | 63 | 6 | 1 | 0 | 20 | 9 |
| M2 | 0 | 4 | 5 | 10 | 35 | 210 | 3 | 67 | 15 | 18 | 1 | 0 | 20 | 11 |
| M3 | 0 | 7 | 5 | 10 | 35 | 225 | 5 | 77 | 8 | 13 | 2 | 1 | 20 | 10 |
| M4 | 0 | 6 | 6 | 13 | 46 | 374 | 3 | 55 | 23 | 18 | 2 | 1 | 21 | 10 |
| M5 | 0 | 3 | 6 | 12 | 44 | 321 | 2 | 70 | 24 | 4 | 2 | 1 | 20 | 9 |
| C1 | 0 | 7 | 5 | 12 | 42 | 312 | 3 | 46 | 1 | 51 | 2 | 1 | 23 | 11 |
| M1 | 3 | 7 | 5 | 11 | 33 | 250 | 4 | 66 | 29 | 4 | 1 | 0 | 23 | 9 |
| M2 | 3 | 6 | 5 | 11 | 35 | 182 | 4 | 67 | 20 | 11 | 1 | 0 | 21 | 10 |
| M3 | 3 | 9 | 6 | 12 | 42 | 122 | 6 | 66 | 25 | 8 | 1 | 0 | 19 | 11 |
| M4 | 3 | 10 | 5 | 12 | 40 | 345 | 2 | 20 | 1 | 77 | 2 | 1 | 22 | 9 |
| M5 | 3 | 6 | 4 | 11 | 31 | 248 | 1 | 24 | 40 | 34 | 2 | 1 | 26 | 8 |
| C1 | 3 | 15 | 5 | 12 | 45 | 258 | 3 | 18 | 1 | 79 | 2 | 1 | 22 | 11 |
| M1 | 6 | 5 | 5 | 11 | 37 | 256 | 4 | 82 | 14 | 3 | 1 | 0 | 20 | 8 |
| M2 | 6 | 5 | 5 | 10 | 37 | 189 | 3 | 65 | 15 | 18 | 2 | 0 | 20 | 11 |
| M3 | 6 | 6 | 5 | 10 | 35 | 234 | 5 | 84 | 13 | 1 | 2 | 1 | 20 | 9 |
| M4 | 6 | 6 | 6 | 12 | 41 | 333 | 3 | 53 | 29 | 16 | 1 | 1 | 22 | 9 |
| M5 | 6 | 2 | 6 | 11 | 42 | 277 | 2 | 75 | 17 | 7 | 1 | 1 | 20 | 8 |
| C1 | 6 | 3 | 5 | 12 | 44 | 23 | 1 | 43 | 1 | 48 | 7 | 0 | 22 | 10 |
| M1 | 9 | 6 | 5 | 11 | 37 | 300 | 5 | 75 | 17 | 6 | 1 | 0 | 20 | 9 |
| M2 | 9 | 7 | 5 | 10 | 34 | 213 | 4 | 57 | 19 | 21 | 2 | 0 | 20 | 10 |
| M3 | 9 | 8 | 5 | 11 | 38 | 252 | 6 | 82 | 5 | 10 | 2 | 1 | 20 | 10 |
| M4 | 9 | 8 | 6 | 12 | 41 | 332 | 4 | 51 | 1 | 46 | 1 | 1 | 22 | 9 |
| M5 | 9 | 5 | 5 | 11 | 38 | 329 | 4 | 84 | 7 | 9 | 1 | 0 | 21 | 8 |
| M1 | 12 | 7 | 5 | 11 | 38 | 288 | 6 | 84 | 9 | 3 | 3 | 1 | 20 | 9 |
| M2 | 12 | 7 | 5 | 10 | 32 | 229 | 5 | 79 | 6 | 13 | 2 | 1 | 20 | 11 |
| M3 | 12 | 7 | 5 | 11 | 36 | 282 | 6 | 84 | 5 | 7 | 3 | 1 | 20 | 10 |
| M4 | 12 | 8 | 4 | 12 | 40 | 257 | 3 | 39 | 1 | 60 | 2 | 1 | 30 | 9 |
| M5 | 12 | 4 | 5 | 10 | 35 | 351 | 3 | 87 | 5 | 6 | 2 | 1 | 21 | 8 |
| M1 | 15 | 7 | 5 | 11 | 40 | 292 | 5 | 80 | 7 | 8 | 4 | 1 | 20 | 9 |
| M2 | 15 | 5 | 4 | 9 | 33 | 45 | 3 | 60 | 28 | 9 | 3 | 0 | 19 | 10 |
| M3 | 15 | 7 | 5 | 11 | 35 | 268 | 6 | 79 | 11 | 8 | 2 | 1 | 20 | 9 |
| M4 | 15 | 6 | 5 | 12 | 39 | 389 | 3 | 48 | 1 | 48 | 2 | 1 | 22 | 9 |
| M5 | 15 | 5 | 5 | 10 | 35 | 388 | 4 | 82 | 4 | 13 | 1 | 1 | 21 | 8 |
| M1 | 21 | 6 | 5 | 10 | 35 | 255 | 3 | 55 | 31 | 11 | 2 | 1 | 21 | 9 |
| M2 | 21 | 5 | 5 | 9 | 34 | 251 | 4 | 88 | 3 | 9 | 0 | 0 | 19 | 10 |
| M3 | 21 | 6 | 5 | 11 | 36 | 310 | 5 | 80 | 7 | 11 | 2 | 1 | 20 | 9 |
| M4 | 21 | 6 | 5 | 11 | 40 | 405 | 4 | 66 | 25 | 7 | 2 | 1 | 22 | 9 |
| M5 | 21 | 3 | 5 | 11 | 36 | 330 | 2 | 80 | 13 | 5 | 2 | 1 | 21 | 8 |
| M1 | 28 | 5 | 5 | 10 | 37 | 289 | 2 | 44 | 51 | 2 | 2 | 1 | 20 | 9 |
| M2 | 28 | 3 | 4 | 9 | 28 | 199 | 2 | 67 | 13 | 18 | 2 | 0 | 24 | 10 |
| M3 | 28 | 6 | 5 | 11 | 38 | 255 | 5 | 82 | 8 | 9 | 1 | 0 | 20 | 10 |
| M4 | 28 | 5 | 6 | 13 | 43 | 327 | 2 | 36 | 2 | 61 | 1 | 0 | 23 | 10 |
| M5 | 28 | 3 | 5 | 11 | 36 | 305 | 2 | 60 | 22 | 16 | 2 | 1 | 21 | 8 |

WBC, white blood cells; RBC, red blood cells; HgB, hemoglobin; HCT, hematocrit level; PLT, platelets; Lymph, lymphocytes; Mono., monocytes; Neutr., neutrophils; Eosin., eosinophils; Baso., basophils; MCH, mean corpuscular hemoglobin; MPV, mean platelet volume. M1-M5 are designated identifications of treated NHPs, and C1 is an untreated control NHP.

**Table S2. Hematology measurements from rEBOV-515 + rEBOV-442 cocktail-treated or -untreated individual NHP that were challenged with BDBV. Related to Figure 4**

| Animal ID | Day after challenge | WBC per $mm^3$ ( x $10^3$) | RBC per $mm^3$ (x $10^3$) | HgB (g/dL) | Hct (%) | Platelets per $mm^3$ (x $10^3$ | Lymph per $mm^3$ (x $10^3$) | Lymph. (%) | Mono. per $mm^3$ (x $10^3$) | Neutr. (%) | Eosin. (%) | Baso. (%) | MCH (pg/cell) | MPV (fL/cell) |
|---|---|---|---|---|---|---|---|---|---|---|---|---|---|---|
| M1 | 0 | 13 | 7 | 14 | 47 | 316 | 8 | 58 | 0 | 30 | 9 | 2 | 21 | 9 |
| M2 | 0 | 7 | 6 | 12 | 41 | 279 | 4 | 59 | 0 | 31 | 6 | 2 | 20 | 8 |
| M3 | 0 | 15 | 6 | 13 | 43 | 366 | 9 | 59 | 0 | 34 | 5 | 2 | 22 | 9 |
| M4 | 0 | 11 | 6 | 13 | 43 | 439 | 4 | 39 | 0 | 53 | 6 | 2 | 21 | 8 |
| M5 | 0 | 9 | 7 | 13 | 44 | 297 | 5 | 58 | 0 | 32 | 7 | 2 | 20 | 9 |
| C1 | 0 | 10 | 5 | 12 | 41 | 277 | 7 | 64 | 0 | 30 | 4 | 1 | 22 | 9 |
| M1 | 3 | 13 | 6 | 13 | 43 | 373 | 8 | 62 | 0 | 32 | 4 | 2 | 21 | 9 |
| M2 | 3 | 7 | 6 | 12 | 40 | 309 | 4 | 51 | 0 | 45 | 2 | 1 | 20 | 9 |
| M3 | 3 | 11 | 6 | 13 | 45 | 200 | 7 | 66 | 0 | 26 | 5 | 1 | 21 | 10 |
| M4 | 3 | 14 | 6 | 12 | 42 | 417 | 3 | 25 | 0 | 67 | 6 | 2 | 20 | 9 |
| M5 | 3 | 9 | 7 | 13 | 44 | 294 | 4 | 45 | 0 | 51 | 3 | 1 | 19 | 9 |
| C1 | 3 | 13 | 5 | 11 | 39 | 299 | 7 | 49 | 0 | 47 | 2 | 1 | 21 | 10 |
| M1 | 6 | 10 | 6 | 13 | 46 | 151 | 2 | 21 | 0 | 75 | 3 | 1 | 21 | 9 |
| M2 | 6 | 6 | 5 | 11 | 36 | 238 | 2 | 38 | 0 | 57 | 2 | 1 | 20 | 8 |
| M3 | 6 | 10 | 6 | 13 | 41 | 322 | 5 | 49 | 0 | 48 | 2 | 1 | 22 | 9 |
| M4 | 6 | 10 | 6 | 12 | 44 | 161 | 3 | 31 | 0 | 67 | 1 | 0 | 19 | 9 |
| M5 | 6 | 12 | 6 | 13 | 42 | 168 | 2 | 16 | 0 | 80 | 3 | 1 | 20 | 9 |
| C1 | 6 | 13 | 5 | 12 | 38 | 196 | 2 | 16 | 0 | 82 | 1 | 0 | 23 | 10 |
| M1 | 9 | 15 | 6 | 14 | 47 | 169 | 5 | 34 | 0 | 60 | 4 | 1 | 22 | 10 |
| M2 | 9 | 9 | 5 | 11 | 38 | 195 | 4 | 45 | 0 | 48 | 2 | 1 | 21 | 9 |
| M3 | 9 | 10 | 5 | 12 | 40 | 395 | 7 | 69 | 1 | 17 | 5 | 2 | 23 | 9 |
| M4 | 9 | 11 | 6 | 12 | 42 | 234 | 4 | 37 | 0 | 57 | 2 | 1 | 21 | 11 |
| M5 | 9 | 12 | 6 | 12 | 41 | 108 | 6 | 51 | 0 | 46 | 1 | 0 | 20 | 11 |
| C1 | 9 | 11 | 5 | 12 | 4 | 176 | 2 | 22 | 0 | 76 | 1 | 0 | 23 | 12 |
| M1 | 12 | 19 | 5 | 12 | 41 | 347 | 14 | 74 | 1 | 15 | 3 | 1 | 22 | 10 |
| M2 | 12 | 9 | 5 | 11 | 38 | 302 | 5 | 57 | 0 | 34 | 6 | 2 | 21 | 8 |
| M3 | 12 | 14 | 5 | 12 | 40 | 640 | 10 | 69 | 0 | 27 | 1 | 1 | 23 | 9 |
| M4 | 12 | 26 | 5 | 11 | 39 | 322 | 14 | 55 | 0 | 42 | 1 | 0 | 21 | 10 |
| M5 | 12 | 17 | 5 | 11 | 35 | 282 | 11 | 65 | 2 | 20 | 3 | 1 | 21 | 10 |
| C1 | 12 | 26 | 5 | 12 | 39 | 113 | 5 | 17 | 0 | 79 | 3 | 0 | 23 | 12 |
| C1 | 14 | 23 | 5 | 12 | 40 | 110 | 4 | 17 | 0 | 80 | 2 | 0 | 23 | 11 |
| M1 | 15 | 16 | 5 | 12 | 41 | 442 | 10 | 59 | 0 | 35 | 3 | 2 | 23 | 9 |
| M2 | 15 | 9 | 5 | 10 | 37 | 304 | 5 | 50 | 0 | 46 | 3 | 1 | 20 | 8 |
| M3 | 15 | 18 | 5 | 12 | 39 | 543 | 10 | 54 | 0 | 40 | 4 | 2 | 23 | 8 |
| M4 | 15 | 15 | 5 | 10 | 37 | 477 | 9 | 58 | 0 | 40 | 1 | 1 | 21 | 9 |
| M5 | 15 | 14 | 5 | 11 | 38 | 402 | 6 | 44 | 0 | 51 | 3 | 2 | 21 | 9 |
| M1 | 21 | 16 | 6 | 13 | 45 | 355 | 7 | 48 | 0 | 39 | 10 | 2 | 23 | 9 |
| M2 | 21 | 9 | 5 | 11 | 38 | 330 | 4 | 49 | 0 | 44 | 5 | 2 | 22 | 9 |
| M3 | 21 | 15 | 5 | 11 | 39 | 689 | 7 | 46 | 0 | 50 | 2 | 2 | 21 | 9 |
| M4 | 21 | 16 | 6 | 13 | 43 | 503 | 9 | 58 | 0 | 31 | 8 | 2 | 23 | 9 |
| M5 | 21 | 10 | 5 | 11 | 38 | 434 | 6 | 57 | 0 | 34 | 7 | 2 | 22 | 9 |
| M1 | 28 | 16 | 6 | 14 | 47 | 248 | 10 | 65 | 0 | 24 | 8 | 2 | 23 | 9 |
| M2 | 28 | 9 | 5 | 11 | 39 | 304 | 5 | 61 | 0 | 28 | 8 | 2 | 21 | 8 |
| M3 | 28 | 14 | 5 | 12 | 40 | 441 | 9 | 66 | 0 | 28 | 4 | 2 | 24 | 9 |
| M4 | 28 | 13 | 5 | 11 | 39 | 430 | 7 | 52 | 0 | 41 | 4 | 2 | 21 | 9 |
| M5 | 28 | 11 | 6 | 12 | 41 | 237 | 8 | 69 | 0 | 18 | 12 | 2 | 21 | 10 |
| M1 | 35 | 15 | 6 | 14 | 46 | 329 | 10 | 65 | 0 | 27 | 7 | 2 | 23 | 9 |
| M2 | 35 | 10 | 5 | 11 | 38 | 297 | 6 | 64 | 0 | 29 | 5 | 2 | 22 | 8 |

| | | | | | | | | | | | | | | |
|---|---|---|---|---|---|---|---|---|---|---|---|---|---|---|
| M3 | 35 | 11 | 5 | 12 | 39 | 349 | 8 | 70 | 0 | 26 | 3 | 1 | 23 | 9 |
| M4 | 35 | 14 | 5 | 11 | 38 | 341 | 6 | 46 | 0 | 51 | 1 | 1 | 21 | 8 |
| M5 | 35 | 12 | 5 | 11 | 39 | 199 | 7 | 61 | 0 | 27 | 10 | 2 | 21 | 10 |

Hematology measurement designations are as in **Table S1**. M1-M5 are designated identifications of treated NHPs, and C1 is an untreated control NHP.

#### Table S3. Hematology measurements from rEBOV-515+rEBOV-442 cocktail-treated or -untreated individual NHP that were challenged with SUDV. Related to Figure 4

| Animal ID | Day after challenge | WBC per $mm^3$ ( x $10^3$) | RBC per $mm^3$ (x $10^3$) | HgB (g/dL) | Hct (%) | Platelets per $mm^3$ (x $10^3$ | Lymph per $mm^3$ (x $10^3$) | Lymph. (%) | Mono. per $mm^3$ (x $10^3$) | Neutr. (%) | Eosin. (%) | Baso. (%) | MCH (pg/cell) | MPV (fL/cell) |
|---|---|---|---|---|---|---|---|---|---|---|---|---|---|---|
| C1 | 0 | 5 | 7 | 14 | 49 | 311 | 4 | 81 | 0 | 6 | 3 | 1 | 20 | 9 |
| M1 | 0 | 6 | 6 | 13 | 41 | 265 | 5 | 82 | 0 | 14 | 1 | 0 | 22 | 11 |
| M2 | 0 | 4 | 5 | 12 | 41 | 395 | 3 | 62 | 1 | 20 | 1 | 1 | 22 | 8 |
| M3 | 0 | 6 | 5 | 11 | 37 | 305 | 4 | 75 | 1 | 11 | 1 | 0 | 21 | 8 |
| M4 | 0 | 8 | 4 | 11 | 31 | 183 | 7 | 90 | 0 | 7 | 0 | 0 | 26 | 9 |
| M5 | 0 | 7 | 6 | 12 | 44 | 431 | 3 | 46 | 3 | 12 | 4 | 2 | 22 | 9 |
| C1 | 4 | 16 | 6 | 12 | 46 | 255 | 2 | 11 | 0 | 85 | 2 | 1 | 19 | 8 |
| M1 | 4 | 13 | 5 | 12 | 42 | 241 | 2 | 16 | 5 | 42 | 3 | 1 | 22 | 11 |
| M2 | 4 | 17 | 5 | 11 | 37 | 327 | 1 | 6 | 0 | 89 | 2 | 1 | 22 | 9 |
| M3 | 4 | 13 | 5 | 10 | 34 | 240 | 1 | 11 | 0 | 85 | 3 | 1 | 20 | 8 |
| M4 | 4 | 12 | 5 | 10 | 35 | 189 | 7 | 62 | 4 | 6 | 1 | 0 | 22 | 9 |
| M5 | 4 | 30 | 5 | 12 | 43 | 352 | 1 | 4 | 0 | 92 | 2 | 1 | 22 | 9 |
| C1 | 7 | 5 | 6 | 11 | 41 | 87 | 2 | 28 | 2 | 25 | 3 | 0 | 20 | 10 |
| M1 | 7 | 8 | 5 | 11 | 38 | 270 | 3 | 41 | 3 | 28 | 1 | 1 | 22 | 11 |
| M2 | 7 | 12 | 5 | 10 | 34 | 316 | 2 | 13 | 6 | 37 | 2 | 1 | 21 | 8 |
| M3 | 7 | 10 | 5 | 10 | 38 | 232 | 2 | 17 | 0 | 73 | 7 | 2 | 19 | 8 |
| M4 | 7 | 14 | 5 | 10 | 35 | 256 | 4 | 26 | 8 | 13 | 2 | 1 | 21 | 9 |
| M5 | 7 | 17 | 5 | 11 | 40 | 392 | 2 | 12 | 0 | 83 | 2 | 1 | 21 | 9 |
| C1 | 10 | 3 | 6 | 11 | 40 | 27 | 1 | 27 | 0 | 67 | 5 | 0 | 19 | 10 |
| M1 | 10 | 6 | 5 | 11 | 38 | 445 | 5 | 86 | 1 | 2 | 1 | 0 | 21 | 10 |
| M2 | 10 | 20 | 4 | 10 | 32 | 352 | 3 | 14 | 9 | 41 | 1 | 1 | 22 | 9 |
| M3 | 10 | 6 | 5 | 9 | 33 | 358 | 4 | 71 | 1 | 8 | 2 | 1 | 19 | 8 |
| M4 | 10 | 8 | 4 | 9 | 29 | 330 | 7 | 87 | 1 | 5 | 0 | 0 | 22 | 9 |
| M5 | 10 | 12 | 5 | 10 | 38 | 572 | 4 | 37 | 0 | 58 | 3 | 2 | 21 | 8 |
| M1 | 14 | 6 | 5 | 11 | 38 | 472 | 4 | 77 | 1 | 9 | 1 | 1 | 21 | 10 |
| M2 | 14 | 6 | 4 | 10 | 32 | 659 | 4 | 62 | 0 | 26 | 3 | 2 | 23 | 8 |
| M3 | 14 | 6 | 5 | 10 | 36 | 429 | 4 | 58 | 0 | 36 | 5 | 2 | 19 | 8 |
| M4 | 14 | 7 | 4 | 9 | 35 | 464 | 7 | 94 | 0 | 2 | 0 | 0 | 21 | 9 |
| M5 | 14 | 17 | 4 | 10 | 34 | 423 | 3 | 15 | 9 | 27 | 2 | 1 | 23 | 8 |
| M1 | 21 | 5 | 5 | 12 | 35 | 275 | 5 | 82 | 0 | 6 | 2 | 1 | 25 | 11 |
| M2 | 21 | 6 | 5 | 10 | 37 | 441 | 2 | 38 | 0 | 56 | 5 | 2 | 22 | 8 |
| M3 | 21 | 5 | 5 | 10 | 37 | 367 | 4 | 70 | 0 | 20 | 4 | 2 | 19 | 8 |
| M4 | 21 | 8 | 5 | 11 | 37 | 397 | 7 | 91 | 0 | 5 | 1 | 1 | 23 | 9 |
| M5 | 21 | 8 | 5 | 11 | 37 | 432 | 2 | 30 | 4 | 17 | 2 | 1 | 23 | 9 |
| M1 | 28 | 5 | 6 | 12 | 43 | 223 | 4 | 80 | 0 | 9 | 2 | 1 | 20 | 11 |
| M2 | 28 | 4 | 5 | 11 | 40 | 365 | 2 | 49 | 0 | 47 | 3 | 1 | 22 | 8 |
| M3 | 28 | 4 | 5 | 10 | 34 | 167 | 3 | 79 | 1 | 2 | 2 | 1 | 20 | 9 |
| M4 | 28 | 9 | 5 | 10 | 35 | 168 | 8 | 91 | 0 | 2 | 1 | 0 | 22 | 10 |
| M5 | 28 | 7 | 5 | 10 | 36 | 295 | 3 | 42 | 3 | 12 | 3 | 1 | 22 | 9 |

Hematology measurement designations are as in **Table S1**. M1-M5 are designated identifications of treated NHPs, and C1 is an untreated control NHP.

**Table S4. Blood biochemistry measurements from rEBOV-515 + rEBOV-442 cocktail-treated or -untreated NHP that were challenged with EBOV. Related to Figure 4**

| **Animal ID** | **Day after challenge** | **Glucose** (mg/dL) | **BUN** (mg/dL) | **Creatinine** (mg/dL) | **Albumin** (g/dL) | **TP** (g/dL) | **ALT** (U/L) | **AST** (U/L) | **ALP** (U/L) | **GGT** (U/L) | **AMY** (U/L) | **C-reactive protein** (mg/dL) |
|---|---|---|---|---|---|---|---|---|---|---|---|---|
| M1 | 0 | 88 | 34 | 1 | 4 | 8 | 46 | 32 | 252 | 72 | 273 | <5.0 |
| M2 | 0 | 98 | 20 | 1 | 4 | 7 | 58 | 26 | 204 | 63 | 256 | <5.0 |
| M3 | 0 | 76 | 17 | 0 | 4 | 7 | 45 | 32 | 440 | 104 | 166 | <5.0 |
| M4 | 0 | 65 | 20 | 1 | 3 | 7 | 123 | 60 | 209 | 78 | 307 | <5.0 |
| M5 | 0 | 79 | 14 | 1 | 3 | 6 | 38 | 32 | 311 | 77 | 320 | <5.0 |
| C1 | 0 | 93 | 15 | 0 | 3 | 8 | 41 | 27 | 266 | 92 | 296 | <5.0 |
| M1 | 3 | 94 | 43 | 1 | 3 | 8 | 48 | 32 | 255 | 72 | 297 | <5.0 |
| M2 | 3 | 82 | 21 | 1 | 3 | 7 | 62 | 29 | 170 | 59 | 276 | <5.0 |
| M3 | 3 | 85 | 23 | 1 | 4 | 7 | 44 | 31 | 414 | 100 | 203 | <5.0 |
| M4 | 3 | 67 | 20 | 1 | 3 | 6 | 147 | 67 | 196 | 75 | 324 | <5.0 |
| M5 | 3 | 61 | 20 | 1 | 3 | 6 | 33 | 32 | 310 | 72 | 370 | 5.0 |
| C1 | 3 | 92 | 14 | 1 | 3 | 8 | 41 | 29 | 230 | 82 | 335 | 15.8 |
| M1 | 6 | 85 | 38 | 1 | 3 | 7 | 38 | 30 | 216 | 69 | 353 | <5.0 |
| M2 | 6 | 69 | 22 | 1 | 3 | 7 | 51 | 32 | 146 | 58 | 259 | <5.0 |
| M3 | 6 | 74 | 21 | 1 | 3 | 7 | 35 | 35 | 336 | 94 | 172 | <5.0 |
| M4 | 6 | 72 | 25 | 1 | 3 | 7 | 102 | 48 | 182 | 75 | 301 | <5.0 |
| M5 | 6 | 62 | 23 | 1 | 3 | 6 | 47 | 63 | 301 | 66 | 316 | <5.0 |
| C1 | 6 | 13 | 101 | 6 | 2 | 7 | 931 | 4170* | 1708 | 484 | 514 | 160.0 |
| M1 | 9 | 82 | 28 | 1 | 3 | 8 | 40 | 33 | 205 | 66 | 250 | <5.0 |
| M2 | 9 | 73 | 19 | 1 | 3 | 7 | 55 | 32 | 142 | 57 | 232 | <5.0 |
| M3 | 9 | 83 | 23 | 1 | 3 | 7 | 38 | 38 | 337 | 89 | 210 | <5.0 |
| M4 | 9 | 69 | 24 | 1 | 3 | 7 | 103 | 60 | 177 | 73 | 302 | <5.0 |
| M5 | 9 | 63 | 26 | 1 | 3 | 6 | 48 | 46 | 271 | 66 | 289 | <5.0 |
| M1 | 12 | 81 | 27 | 1 | 3 | 8 | 34 | 33 | 211 | 68 | 280 | <5.0 |
| M2 | 12 | 84 | 21 | 1 | 3 | 7 | 49 | 30 | 144 | 58 | 278 | <5.0 |
| M3 | 12 | 67 | 23 | 1 | 3 | 7 | 38 | 34 | 360 | 88 | 201 | <5.0 |
| M4 | 12 | 49 | 23 | 1 | 3 | 7 | 131 | 70 | 167 | 72 | 320 | <5.0 |
| M5 | 12 | 62 | 22 | 1 | 3 | 6 | 34 | 37 | 274 | 65 | 316 | <5.0 |
| M1 | 15 | 89 | 25 | 1 | 3 | 8 | 32 | 31 | 214 | 66 | 260 | 8.3 |
| M2 | 15 | 81 | 19 | 1 | 3 | 7 | 63 | 38 | 138 | 56 | 249 | <5.0 |
| M3 | 15 | 71 | 19 | 1 | 3 | 7 | 44 | 35 | 384 | 87 | 196 | <5.0 |
| M4 | 15 | 61 | 21 | 1 | 3 | 7 | 124 | 54 | 168 | 69 | 300 | <5.0 |
| M5 | 15 | 74 | 17 | 1 | 3 | 6 | 31 | 31 | 292 | 66 | 341 | <5.0 |
| M1 | 21 | 67 | 34 | 0 | 3 | 8 | 29 | 29 | 203 | 64 | 263 | <5.0 |
| M2 | 21 | 84 | 19 | 1 | 3 | 7 | 50 | 28 | 108 | 57 | 254 | <5.0 |
| M3 | 21 | 72 | 21 | 1 | 4 | 7 | 40 | 37 | 366 | 91 | 186 | <5.0 |
| M4 | 21 | 67 | 24 | 1 | 3 | 7 | 85 | 42 | 147 | 63 | 310 | <5.0 |
| M5 | 21 | 61 | 24 | 1 | 3 | 6 | 26 | 27 | 316 | 68 | 358 | <5.0 |
| M1 | 28 | 58 | 32 | 1 | 3 | 7 | 32 | 29 | 208 | 64 | 240 | 5.8 |
| M2 | 28 | 74 | 20 | 1 | 4 | 7 | 74 | 36 | 120 | 58 | 257 | <5.0 |
| M3 | 28 | 66 | 22 | 1 | 4 | 7 | 38 | 35 | 362 | 93 | 166 | <5.0 |
| M4 | 28 | 45 | 23 | 1 | 3 | 7 | 81 | 42 | 129 | 55 | 275 | <5.0 |
| M5 | 28 | 53 | 24 | 1 | 3 | 5 | 22 | 39 | 243 | 57 | 272 | <5.0 |

BUN, blood urea nitrogen; TP, total protein; ALT, alanine aminotransferase; AST, aspartate aminotransferase; ALP, alkaline phosphatase; GGT, gamma-glutamyl transpeptidase; AMY, amylase. * Sample was assessed at 10-fold dilution, and the value from the measurement was multiplied by ten to generate the recorded value. M1-M5 are designated identifications of treated NHPs, and C1 is an untreated control NHP.

**Table S5. Blood biochemistry measurements from rEBOV-515+rEBOV-442 cocktail-treated or -untreated NHP that were challenged with BDBV. Related to Figure 4**

| Animal ID | Day after challenge | Glucose (mg/dL) | BUN (mg/dL) | Creatinine (mg/dL) | Albumin (g/dL) | TP (g/dL) | ALT (U/L) | AST (U/L) | ALP (U/L) | GGT (U/L) | AMY (U/L) | C-reactive protein (mg/dL) |
|---|---|---|---|---|---|---|---|---|---|---|---|---|
| M1 | 0 | 87 | 19 | 1 | 3 | 7 | 31 | 33 | 463 | 63 | 338 | <5.0 |
| M2 | 0 | 64 | 19 | 1 | 3 | 7 | 28 | 47 | 294 | 73 | 270 | 9 |
| M3 | 0 | 84 | 21 | 1 | 3 | 7 | 27 | 31 | 280 | 66 | 467 | 9 |
| M4 | 0 | 83 | 20 | 1 | 3 | 7 | 55 | 35 | 222 | 73 | 458 | <5.0 |
| M5 | 0 | 93 | 20 | 1 | 3 | 7 | 30 | 40 | 270 | 46 | 296 | 7 |
| C1 | 0 | 89 | 18 | 1 | 3 | 6 | 27 | 30 | 171 | 38 | 217 | <5.0 |
| M1 | 3 | 67 | 19 | 1 | 3 | 7 | 37 | 33 | 474 | 60 | 324 | <5.0 |
| M2 | 3 | 66 | 19 | 1 | 3 | 8 | 22 | 40 | 309 | 74 | 261 | 7 |
| M3 | 3 | 85 | 20 | 1 | 3 | 7 | 35 | 39 | 290 | 63 | 487 | 11 |
| M4 | 3 | 78 | 21 | 1 | 3 | 7 | 65 | 34 | 216 | 75 | 462 | <5.0 |
| M5 | 3 | 76 | 20 | 0 | 3 | 8 | 40 | 41 | 278 | 16 | 291 | 7 |
| C1 | 3 | 90 | 17 | 1 | 3 | 6 | 33 | 29 | 166 | 38 | 207 | <5.0 |
| M1 | 6 | 87 | 20 | 1 | 3 | 7 | 33 | 55 | 387 | 53 | 265 | 123 |
| M2 | 6 | 71 | 18 | 1 | 3 | 7 | 22 | 44 | 286 | 66 | 253 | 7 |
| M3 | 6 | 82 | 18 | 1 | 3 | 7 | 32 | 29 | 247 | 54 | 336 | 106 |
| M4 | 6 | 100 | 15 | 1 | 3 | 7 | 29 | 52 | 171 | 34 | 178 | 101 |
| M5 | 6 | 83 | 20 | 1 | 3 | 7 | 30 | 44 | 216 | 37 | 240 | 136 |
| C1 | 6 | 93 | 17 | 1 | 2 | 7 | 57 | 106 | 294 | 81 | 336 | 164 |
| M1 | 9 | 76 | 22 | 1 | 3 | 8 | 168 | 242 | 403 | 73 | 232 | 25 |
| M2 | 9 | 66 | 26 | 1 | 3 | 7 | 26 | 52 | 312 | 66 | 267 | 14 |
| M3 | 9 | 82 | 23 | 1 | 3 | 7 | 37 | 29 | 218 | 55 | 456 | 26 |
| M4 | 9 | 80 | 26 | 1 | 2 | 7 | 147 | 66 | 421 | 120 | 303 | 33 |
| M5 | 9 | 78 | 22 | 1 | 3 | 7 | 116 | 94 | 273 | 76 | 196 | 39 |
| C1 | 9 | 81 | 33 | 1 | 2 | 6 | 55 | 114 | 471 | 186 | 150 | 23 |
| M1 | 12 | 74 | 17 | 1 | 3 | 7 | 82 | 42 | 297 | 63 | 344 | 7 |
| M2 | 12 | 66 | 19 | 1 | 3 | 7 | 29 | 46 | 379 | 83 | 287 | 10 |
| M3 | 12 | 90 | 19 | 1 | 3 | 7 | 31 | 30 | 143 | 61 | 528 | 11 |
| M4 | 12 | 81 | 22 | 1 | 2 | 7 | 56 | 38 | 315 | 98 | 339 | 7 |
| M5 | 12 | 72 | 23 | 1 | 3 | 7 | 51 | 47 | 202 | 52 | 356 | 9 |
| C1 | 12 | 107 | >180 | 6 | 2 | 6 | 66 | 164 | 566 | 338 | 305 | <5.0 |
| C1 | 14 | 190 | 380* | 12 | 1 | 5 | 85 | 227 | 357 | 198 | 63 | <5.0 |
| M1 | 15 | 66 | 20 | 1 | 3 | 7 | 49 | 34 | 340 | 64 | 337 | 6 |
| M2 | 15 | 55 | 20 | 1 | 3 | 7 | 34 | 50 | 309 | 73 | 245 | 10 |
| M3 | 15 | 75 | 21 | 1 | 3 | 7 | 35 | 47 | 249 | 61 | 398 | 22 |
| M4 | 15 | 72 | 25 | 1 | 3 | 7 | 68 | 47 | 309 | 108 | 414 | 6 |
| M5 | 15 | 64 | 21 | 1 | 3 | 7 | 41 | 48 | 205 | 49 | 329 | 9 |
| M1 | 21 | 76 | 17 | 0 | 3 | 7 | 44 | 35 | 418 | 56 | 340 | 5 |
| M2 | 21 | 70 | 18 | 1 | 3 | 7 | 28 | 43 | 273 | 74 | 259 | 8 |
| M3 | 21 | 75 | 24 | 1 | 3 | 7 | 56 | 32 | 245 | 97 | 414 | <5.0 |
| M4 | 21 | 83 | 20 | 1 | 3 | 7 | 25 | 29 | 272 | 63 | 415 | 9 |
| M5 | 21 | 77 | 20 | 1 | 3 | 8 | 27 | 41 | 207 | 42 | 281 | 7 |
| M1 | 28 | 85 | 22 | 1 | 3 | 7 | 28 | 37 | 271 | 64 | 418 | 8 |
| M2 | 28 | 66 | 19 | 1 | 3 | 7 | 23 | 45 | 272 | 69 | 259 | 6 |
| M3 | 28 | 85 | 20 | 1 | 3 | 7 | 39 | 33 | 379 | 52 | 364 | <5.0 |
| M4 | 28 | 71 | 24 | 1 | 3 | 7 | 56 | 40 | 214 | 75 | 431 | <5.0 |
| M5 | 28 | 73 | 21 | 1 | 3 | 7 | 29 | 41 | 202 | 40 | 311 | 7 |
| M1 | 35 | 79 | 19 | 1 | 3 | 7 | 53 | 34 | 395 | 53 | 365 | <5.0 |
| M2 | 35 | 63 | 17 | 1 | 3 | 7 | 22 | 41 | 275 | 66 | 288 | 8 |
| M3 | 35 | 73 | 21 | 1 | 3 | 6 | 33 | 31 | 283 | 60 | 396 | 11 |
| M4 | 35 | 78 | 25 | 1 | 3 | 6 | 61 | 43 | 208 | 64 | 40 | 5 |

| | | | | | | | | | | | | |
|---|---|---|---|---|---|---|---|---|---|---|---|---|
| M5 | 35 | 66 | 21 | 1 | 3 | 6 | 35 | 36 | 231 | 42 | 369 | 6 |

Blood chemistry measurement designations are as in **Table S4**. * Sample was assessed at 5-fold dilution, and the value from the measurement was multiplied by five to generate the recorded value. M1-M5 are designated identifications of treated NHPs, and C1 is an untreated control NHP.

#### Table S6. Blood biochemistry measurements from rEBOV-515+rEBOV-442 cocktail-treated or -untreated NHP that were challenged with SUDV. Related to Figure 4

| Animal ID | Day after challenge | Glucose (mg/dL) | BUN (mg/dL) | Creatinine (mg/dL) | Albumin (g/dL) | TP (g/dL) | ALT (U/L) | AST (U/L) | ALP (U/L) | GGT (U/L) | AMY (U/L) | C-reactive protein (mg/dL) |
|---|---|---|---|---|---|---|---|---|---|---|---|---|
| C1 | 0 | 97 | 23 | 1 | 3 | 7 | 119 | 43 | 388 | 82 | 397 | <5.0 |
| M1 | 0 | 115 | 27 | 1 | 4 | 7 | 64 | 35 | 262 | 81 | 345 | <5.0 |
| M2 | 0 | 82 | 15 | 1 | 3 | 7 | 84 | 33 | 247 | 73 | 286 | <5.0 |
| M3 | 0 | 83 | 19 | 1 | 3 | 7 | 67 | 33 | 217 | 59 | 311 | <5.0 |
| M4 | 0 | 89 | 28 | 1 | 3 | 7 | 49 | 35 | 213 | 67 | 349 | <5.0 |
| M5 | 0 | 80 | 23 | 1 | 3 | 7 | 34 | 32 | 258 | 61 | 344 | <5.0 |
| C1 | 4 | 89 | 20 | 1 | 3 | 7 | 82 | 38 | 322 | 71 | 211 | 125 |
| M1 | 4 | 88 | 17 | 1 | 3 | 6 | 61 | 31 | 202 | 55 | 228 | 109 |
| M2 | 4 | 93 | 14 | 0 | 3 | 7 | 76 | 34 | 208 | 65 | 220 | 103 |
| M3 | 4 | 88 | 19 | 1 | 3 | 7 | 50 | 33 | 226 | 67 | 228 | 129 |
| M4 | 4 | 81 | 24 | 1 | 3 | 6 | 50 | 34 | 169 | 60 | 393 | <5.0 |
| M5 | 4 | 86 | 17 | 1 | 3 | 7 | 44 | 34 | 241 | 54 | 226 | 139 |
| C1 | 7 | 73 | 24 | 1 | 2 | 6 | 298 | 379 | 990 | 123 | 162 | 121 |
| M1 | 7 | 63 | 20 | 1 | 3 | 7 | 73 | 64 | 230 | 50 | 269 | 6 |
| M2 | 7 | 80 | 13 | 1 | 3 | 6 | 59 | 31 | 199 | 55 | 180 | 29 |
| M3 | 7 | 75 | 23 | 1 | 3 | 7 | 53 | 34 | 248 | 64 | 227 | 26 |
| M4 | 7 | 81 | 29 | 1 | 3 | 7 | 42 | 34 | 172 | 56 | 293 | 21 |
| M5 | 7 | 82 | 21 | 1 | 3 | 7 | 87 | 39 | 391 | 50 | 219 | 90 |
| C1 | 10 | 59 | 36 | 1 | 2 | 6 | 159 | 478 | 1364 | 165 | 124 | 140 |
| M1 | 10 | 56 | 17 | 1 | 3 | 7 | 53 | 32 | 208 | 46 | 269 | <5.0 |
| M2 | 10 | 108 | 16 | 1 | 2 | 6 | 28 | 30 | 308 | 51 | 230 | 168 |
| M3 | 10 | 71 | 19 | 1 | 3 | 7 | 37 | 30 | 225 | 61 | 280 | <5.0 |
| M4 | 10 | 78 | 23 | 0 | 3 | 6 | 43 | 35 | 174 | 49 | 291 | 17 |
| M5 | 10 | 72 | 21 | <0.2 | 2 | 7 | 55 | 33 | 290 | 49 | 276 | <5.0 |
| M1 | 14 | 67 | 18 | 0 | 3 | 7 | 54 | 29 | 228 | 53 | 287 | <5.0 |
| M2 | 14 | 55 | 13 | 0 | 2 | 8 | 33 | 30 | 310 | 61 | 198 | 6 |
| M3 | 14 | 69 | 20 | 1 | 3 | 7 | 31 | 32 | 231 | 66 | 273 | <5.0 |
| M4 | 14 | 74 | 23 | 1 | 3 | 7 | 34 | 29 | 174 | 52 | 321 | <5.0 |
| M5 | 14 | 73 | 20 | 0 | 3 | 7 | 38 | 34 | 248 | 51 | 285 | <5.0 |
| M1 | 21 | 61 | 19 | 1 | 3 | 7 | 66 | 36 | 231 | 56 | 300 | <5.0 |
| M2 | 21 | 76 | 15 | 1 | 3 | 7 | 38 | 33 | 233 | 66 | 265 | <5.0 |
| M3 | 21 | 79 | 24 | 1 | 3 | 7 | 32 | 27 | 230 | 68 | 313 | <5.0 |
| M4 | 21 | 72 | 26 | 1 | 3 | 7 | 38 | 31 | 160 | 54 | 356 | <5.0 |
| M5 | 21 | 76 | 24 | 0 | 3 | 7 | 36 | 32 | 234 | 54 | 313 | <5.0 |
| M1 | 28 | 59 | 20 | 1 | 3 | 7 | 76 | 34 | 235 | 61 | 294 | <5.0 |
| M2 | 28 | 70 | 16 | 1 | 3 | 7 | 41 | 27 | 213 | 57 | 266 | <5.0 |
| M3 | 28 | 70 | 23 | 1 | 4 | 7 | 37 | 31 | 212 | 67 | 296 | <5.0 |
| M4 | 28 | 66 | 29 | 1 | 3 | 7 | 36 | 36 | 169 | 56 | 308 | <5.0 |
| M5 | 28 | 69 | 23 | 1 | 3 | 7 | 28 | 29 | 179 | 50 | 275 | <5.0 |

Hematology measurement designations are as in **Table S1**. M1-M5 are designated identifications of treated NHPs, and C1 is an untreated control NHP.

**Table S7. Cryo-EM data collection and statistics of EBOV GP ΔMucΔTM:rEBOV-515:rEBOV-442 Fab complex. Related to Figure 6**

| Map | EBOV GPΔMucΔTM:rEBOV-442 + rEBOV-515 |
|---|---|
| **Sample vitrification** | - |
| Concentration | 3 mg/mL |
| Grid type | 1.2/1.3 Quantifoil |
| Detergent | 0.01% (w/v) F-OM |
| Blot time | 6 s |
| **Data collection** | 19aug23b |
| Microscope | Titan Krios |
| Voltage (kV) | 300 |
| Detector | K2 Summit |
| Recording mode | Counting |
| Magnification | 48,543 |
| Movie micrograph pixel size (Å) | 1.03 |
| Dose rate (e-/[(camera pixel)*s]) | 6.43 |
| Number of frames per movie | 42 |
| Frame exposure time (ms) | 250 |
| Movie micrograph exposure time (s) | 10.5 |
| Total dose (e-/Å$^2$) | 50 |
| Defocus range (μm) | -1 to -3.5 |
| **EM data processing** | - |
| Software used | Relion 3.1b |
| Number of movie micrographs | 1,020 |
| Number of particles in map | 21,285 |
| Symmetry | C3 |
| Map resolution (FSC 0.143;Å) | 3.3 |
| Map sharpening B-factor (Å$^2$) | -92 |
| **Structure building and validation** | - |
| Number of atoms in deposited model | 18,783 |
| GP1 | 5,541 |
| GP2 | 2,373 |
| rEBOV-442 | 5,571 |
| rEBOV-515 | 5,298 |
| Glycans | 534 |
| MolProbity score | 1.02 |

| | |
|---|---|
| Clashscore | 1.84 |
| EMRinger score | 3.62 |
| RMSD from ideal | - |
| Bond length (Å) | 0.021 |
| Bond angle (degrees) | 1.8 |
| Ramachandran plot | - |
| Favored (%) | 97.65 |
| Allowed (%) | 1.69 |
| Outliers (%) | 0.65 |
| Average B-factor | 128 |
